# Characterisation of an Omnitrap-Orbitrap platform equipped with IRMPD, UVPD and ExD for the analysis of peptides and proteins

**DOI:** 10.1101/2023.05.15.540788

**Authors:** Athanasios Smyrnakis, Nikita Levin, Mariangela Kosmopoulou, Ajay Jha, Kyle Fort, Alexander Makarov, Dimitris Papanastasiou, Shabaz Mohammed

## Abstract

We describe an instrument configuration based on the Orbitrap Exploris 480 mass spectrometer that has been coupled to an Omnitrap platform. The Omnitrap possesses three distinct ion-activation regions, that can be used to perform resonant based collision induced dissociation, several forms of electron associated fragmentation, and ultraviolet photodissociation. Each section can also be combined with infrared multiphoton dissociation. In this work, we demonstrate all these modes of operation on a range of peptides and proteins. The results show that this instrument configuration produces similar data to previous implementations of each activation technique and at similar efficiency levels. We demonstrate that this unique instrument configuration is extremely versatile for the interrogation of polypeptides.

## Introduction

Mass-spectrometric (MS) analysis has achieved remarkable results in the analysis of primary structures of biomolecules.^1^ Until recently, such analysis has mainly relied on collision-induced dissociation (CID), in which ionised molecules of analytes are accelerated or resonantly excited to collide with molecules of buffer gas, which in turn leads to the dissociation of the most labile bonds. In the last few years, mass spectrometer designs have started to move beyond performing simply efficient and speedy CID, expanding their abilities to resolve analytes (e.g., ion mobility^2, 3^) and to provide access to alternative activation techniques.^4, 5^ These expanded abilities are necessary to address the complicated structure of many biomolecules, which is known to determine their biological functions and govern their reaction kinetics. Proteins have been under close focus due to their determining role in life cycles of organisms and the suitability of some to act as therapeutic agents.^6^ Substantial efforts were therefore invested by the MS community in the development of dissociation techniques which would yield exhaustive information about all levels of protein structures and localise and characterise their post translation modifications (PTMs).^5^

In early days of biological mass spectrometry, ion-trap collision-induced dissociation (CID) and beam-type CID were the primary techniques for fragmentation of gaseous ions in quadrupole and linear ion traps (LIT),^7, 8^ whereas in ion cyclotron resonance (ICR) MS, which requires high vacuum incompatible with CID, various fragmentation methods including sustained off-resonance irradiation (SORI) and infrared multiphoton dissociation (IRMPD) were implemented.^9^ As CID, SORI, and IRMPD are typically charge directed and break the weakest bond in a molecule, a comprehensive fragmentation is often prevented, which can be further aggravated if there are labile PTMs such as phosphorylation, sulfation, or glycosylation. The discovery of electron capture dissociation (ECD) by Zubarev and co-workers, in which the recombination of low-energy free electrons with multiply charged precursor results in a gentle cleavage of a C_α_-N bond in a peptide backbone,^10, 11^ allowed for superior localisation of labile PTMs and more robust sequencing of modified peptides, although efficiency requires higher charge states^12^. Due to the incompatibility of free electrons with radiofrequency (RF) based ion traps, the reaction has remained applicable exclusively in ICR MS, even though attempts were made to incorporate ECD into time-of-flight^13^ and Orbitrap™ ^14, 15^ instruments using an electromagnetostatic cell,^16^ into two-dimensional^17^ and three-dimensional ion traps^18^ using weak magnetic fields, and into a digital ion trap without using any magnetic field to focus the electrons.^19^ Hunt and co-workers developed the electron transfer dissociation (ETD) technique as an alternative to ECD for RF-based ion traps.^20^ In this reaction, multiply-charged precursor cations are mixed with negatively charged reagent molecules to facilitate the reaction of electron transfer from anions to cations resulting in dissociation of C_α_-N bonds similar to ECD. The application of ETD has been standardised in Orbitrap hybrid instruments, in which high-resolution Orbitrap mass analyser is coupled with a linear ion trap.^21^ The kinetics of both ECD and ETD is charge-driven making them amenable for top-down analysis of high-charge states of proteins;^14, 15, 22-25^ however charge-reduced species, representing fragments held together by hydrogen bonds, are very often the main reaction products in ECD and ETD due to the non-ergodic nature of these reactions. To increase the number and intensities of fragment ions, the precursor can be activated by low-energy infrared (IR) irradiation concurrently to the reaction of electron capture or transfer. This coactivation disrupts noncovalent bonds of the secondary and tertiary structure thus giving access to otherwise hidden fragmentation sites. Activated-ion ECD and ETD were named AI-ECD^26-31^ and AI-ETD^32-37^ respectively.

In parallel to the development of electron-based fragmentation techniques, the ultraviolet photodissociation (UVPD) of polypeptides was extensively characterised in time-of-flight^38-42^ and more recently in LIT^43, 44^ instruments and proved to be a potent technique for sequencing and characterisation of whole proteins and proteoforms.^45-48^ In contrast to IRMPD, UVPD provides direct excitation of irradiated ions to their dissociative state, which enables extensive fragmentation of amino-acid backbone while preserving most PTMs.^44^ Typically, a single UV laser pulse is sufficient to acquire a near 100% sequence coverage of small proteins with molecular weights below 20k Da.^48^

Recently, a novel ion trap, Omnitrap platform, has been introduced.^49^ This multi-segmented linear ion trap driven by a rectangular waveform generator allows the incorporation of multiple fragmentation techniques within one MS platform, thus enabling multidimensional multiple-stage tandem MS workflows.^49^ In this paper, we characterise the Omnitrap platform coupled with a Thermo Scientific™ Exploris™ 480 Orbitrap mass spectrometer in its application to the sequencing of peptides and proteins in direct-infusion experiments. The Omnitrap has been equipped with an electron gun (for multiple forms of electron reactions), a UV laser and an IR laser.

## Experimental

### Materials

All chemicals as well as peptides (bradykinin, Glu-fibrinopeptide B, and insulin chain B), ubiquitin (bovine), and myoglobin (equine) were purchased from Sigma-Aldrich (Gillingham, Dorset, UK) and used without further purification. Carbonic anhydrase (bovine) was purchased from Sigma-Aldrich (Gillingham, Dorset, UK) and purified using PD-10 desalting columns (Cytiva, Sheffield, UK). Ion optical simulations were performed in SIMION (simion.com), and the results were found to be in good agreement with the thermal^50^ and non-linear^51^ models of the ion density distribution.

### Top-down MS analysis

The analytes were prepared in standard acidified water/acetonitrile solutions. The detailed information about the analytes can be found in Table S1. The experiments were performed on a Thermo Fisher Scientific Exploris Orbitrap 480 mass spectrometer modified with an Omnitrap platform (see below). The Orbitrap instrument was operating constantly in the MS2 mode with HCD kept at 3 V collision potential to facilitate transfer of ions through the HCD cell. The ions were characterised in the Orbitrap analyser with the mass resolution of 30000 for peptides, 120000 for ubiquitin and 480000 for myoglobin and carbonic anhydrase. The injection times were fixed and set to match the AGC targets of 100000 for peptides and one million for proteins. An ArF ExciStar 200 laser (Coherent, Santa Clara, CA) was used as the source of 193 nm UV light. A FireStar ti60 (Synrad, Mukilteo, WA) laser was used as the source of 10.6 µm IR light with the maximum power output of 60 Wt.

### Data analysis

For annotation of peptide fragments, 100 spectra of each peptide were averaged and manually annotated using lists of fragment masses generated by ProteinProspector v6.4.5. For quantitative analysis of fragmentation yields, 60 spectra (carbonic anhydrase and myoglobin) or 100 spectra (peptides and ubiquitin) were averaged and deconvoluted in Freestyle software (Thermo Fisher, San Jose, CA), intensities of fragments peaks were extracted, and the peaks were identified and quantified in MS-TAFI tool^52^. Averaged raw spectra of fragmented ubiquitin and myoglobin^10+,20+^ as well as IRMPD spectra of all proteins and UVPD of [carbonic anhydrase]^20+^ were instead processed and manually revised in an in-house software for top-down analysis yielding identifications of fragments and sequence coverages. The list of the types of fragments used for analysis of spectra acquired in different fragmentation reactions can be found in Supplementary Information (Table S2).

## Results and Discussion

### Instrument configuration

The instrument consists of an Orbitrap Exploris 480 mass spectrometer that has been modified to contain an Omnitrap platform which is connected via two consecutive RF transfer hexapoles. The Omnitrap consists of nine segments, three of which (Q2, Q5 and Q8, see Figure 1) can also act as discrete fragmentation regions. To ensure efficient trapping, nitrogen buffer gas is injected in a pulsed manner via two general valves installed between segments Q1 and Q2. An electron source is installed in Q5, as described previously^49^. The source injects electrons with user-specified energies in the range between ∼0 and 1000 eV. The infrared laser is on-axis allowing fragmentation in any desired region, although the use of a convex lens focuses the IR beam to the fifth segment Q5. Modelling of 500000 charges suggests the radial spread of the ion cloud falls within a diameter of 2 mm ensuring an excellent overlap between the IR laser beam and ion clouds in all segments (Figure S1) including Q5, where the laser beam is marginally smaller than the ion cloud (Figure S2). The UV laser has been installed on Q8 orthogonally to trap axis. The UV laser beam was focused to the centre of segment Q8 where the beam cross-section is smaller than the size of the ion cloud (Figure S3). This modified apparatus can operate as a typical Orbitrap Exploris instrument or have it operate as a source/analyser for the Omnitrap, where precursor ions are transferred into and out of the Omnitrap through the custom-modified back aperture of the HCD cell. The switching between the two modes is accomplished *via* dedicated tune pages of the Exploris control software.

**Figure 1.**
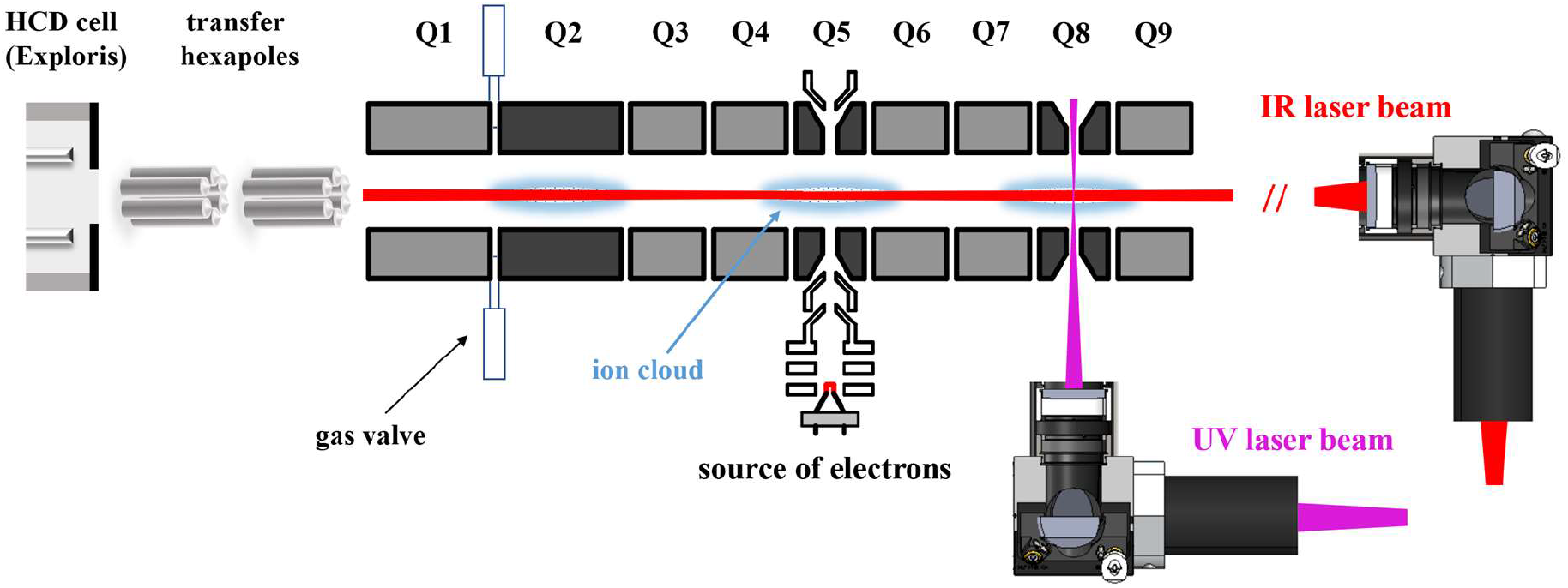
Schematic in-section layout of the Omnitrap platform coupled with the Orbitrap Exploris 480 mass spectrometer.

### Performance of IRMPD

We opted to have the IR laser on-axis so as to allow not only IRMPD but also supplemental activation for the other activation approaches. For characterisation of IRMPD we focused on efficiency, speed, and sensitivity across the three pertinent segments, Q2, Q5 and Q8. Gas pressure, ion trap *q* values, length of IR pulse and power output of the IR laser are known to affect the efficiency^53, 54^. To evaluate the dependence of IRMPD on these parameters, we performed a series of experiments on the model peptide Glu-fibrinopeptide B (EGVNDNEEGFFSAR). The same series of experiments were repeated in segments Q2, Q5, and Q8.

We expected the collisional cooling by the trapping gas to influence fragmentation efficiency of IRMPD. As the temporal profile of pressure in the Omnitrap follows an exponential decay (Figure S4), we expected that increasing the delay between a gas injection and an IR pulse would lead to a more efficient IRMPD. In each of the three segments, the photodissociation efficiency steadily rises until all precursor ions are fragmented when increasing the delay between gas injection and IR triggering (Figure S5). The delay required for each segment was slightly different possibly related to speed of introduction and dissipation of gas for each segment, with typical values of approximately 15 ms. We then investigated the effect of *q* value of the Omnitrap on trapping efficiency of IRMPD. The *q* value defines the depth of potential well and the low-mass cut-off^53^. Increasing *q* value leads to a more efficient depletion of the precursor and generation of fragments (Figure S6). At *q*=0.2 the intensity of the precursor drops to zero and the photodissociation efficiency flattens out. We fixed *q* value at 0.2 in all three segments as a trade-off between low-mass cut-off and efficiency of IRMPD. This value in the Omnitrap is analogous to *q*=0.25 that is used in sinusoidal-RF (*i*.*e*., conventional) ion traps. Overall IRMPD profiles in the Omnitrap are similar to those previously reported for a linear ion trap^54^. Having trapping and timing parameters resolved, we switched to laser pulse duration, and we optimised each segment since laser beam cross section varies across the Omnitrap (Figure S1). As expected, fragmentation increased with increasing pulse length, see Figure S7. Segment Q5 required both shorter pulse lengths and lower laser power to be used while maintaining the same level of fragmentation efficiency as Q2 and Q8 which correlates with our modelling of the size of the laser beam through the Omnitrap (Figure S1). In all experiments described above, the trapping efficiency was initially rising in parallel with increasing photodissociation efficiency until a maximum was reached, after which the trapping efficiency began to decrease due to secondary fragmentation by IR light (Figures S5-S7). After an iterative process of optimising the parameters of IRMPD, we found values that would produce similar spectra in all three segments, Figure S8.

As both high-energy collisional dissociation (HCD) and IRMPD generate typically *b* and *y* fragments, we compared the efficiencies of HCD performed in the HCD cell of Exploris instrument to IRMPD performed in Q5 of the Omnitrap (Figure 2). Both spectra feature near-complete series of *y* fragments, and a few intact *b* fragments are observed exclusively in HCD. The IRMPD spectrum is less congested and is characterised by lower relative intensities in the high mass region compared to its HCD counterpart. The loss of certain fragments is most likely due to secondary fragmentation induced by continuous irradiation by the IR laser. Potentially, the secondary fragmentation process is dictated by cross sectional area of the fragments, larger being more likely to absorb photons.

**Figure 2.**
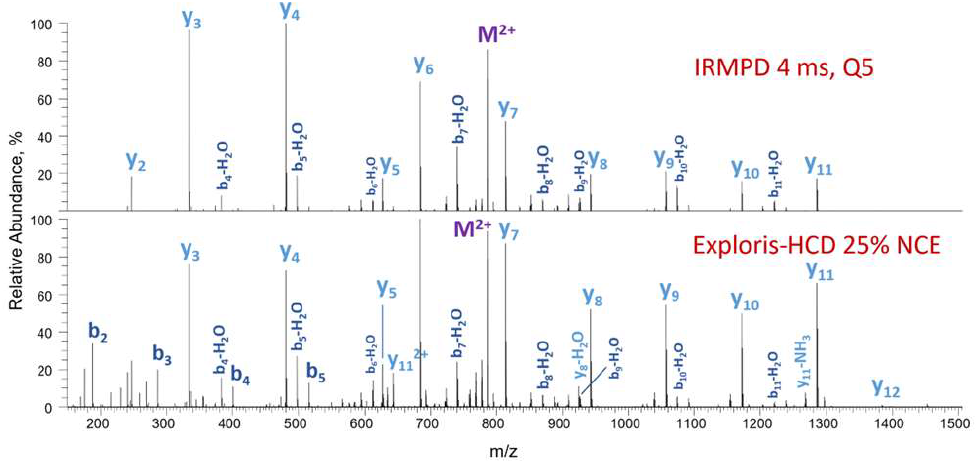
Omnitrap-IRMPD (top) and Exploris-HCD (bottom) spectra of [Glu-fibrinopeptide B]^2+^. IRMPD spectra were acquired after 4 ms of IR irradiation at q=0.2, 30 % laser duty cycle; the IR was triggered 14 ms after the gas pulse. HCD spectra were collected following fragmentation in the HCD cell at 25% of normalised collision energy (NCE).

IRMPD spectra and sequence coverages of ubiquitin^8+^, myoglobin^10+^ and carbonic anhydrase^20+^ acquired in segment Q5 are shown in Figure S9. As the size of the protein increases, the energy needed to induce its fragmentation decreases. In total agreement with earlier results reported for the IRMPD in a linear ion trap^55^, the Omnitrap-IRMPD causes pronounced dissociation of bonds N-terminal to residues of proline and C-terminal to residues of glutamic and aspartic acids, which is reflected in relatively low sequence coverages of the studied proteins (Fig S9).

### Performance of UVPD

We characterised the performance of UVPD in the Omnitrap in a series of experiments with peptides and proteins and assessed if it benefits from supplemental IR-activation. Ions were transferred to segment Q8 and irradiated by pulses of 193 nm UV light with the lasing frequency of 200 Hz. The beam of UV light has an elliptic cross-section and was focused to the centre of segment Q8 using a convex lens (Figure S3). We investigated the number of pulses required for optimal fragmentation and found three pulses (∼10 ms) to be sufficient, above which we started to observe signal loss for peptides and proteins (Figure S10, S11). We noted that efficiency improves as the number of irradiated precursor ions increases which is most likely related to increased density of ion cloud leading to a superior overlap with the laser beam, also reported previously by Fort et al.^56^ As expected, all members of the main fragment ion series are present in the spectra of peptides and proteins along with *a+1, d*, v, *y-1, y-2*, and internal fragments (Figure 3, S12); however, *a+1* and *y* ions are the most abundant types of fragments observed. The distribution of numbers of identified fragments resembles observations by Shaw et al.^48^ as typified by the fragment coverage for ubiquitin^8+^ (Figure S13). The total sequence coverage of ubiquitin^8+^, myoglobin^10+^ and carbonic anhydrase^20+^ in our experiments are 93%, 78% and 64% respectively (Figure S14). These values are broadly in line with those observed by Brodbelt and co-workers who utilise the same laser.^48, 57, 58^

**Figure 3.**
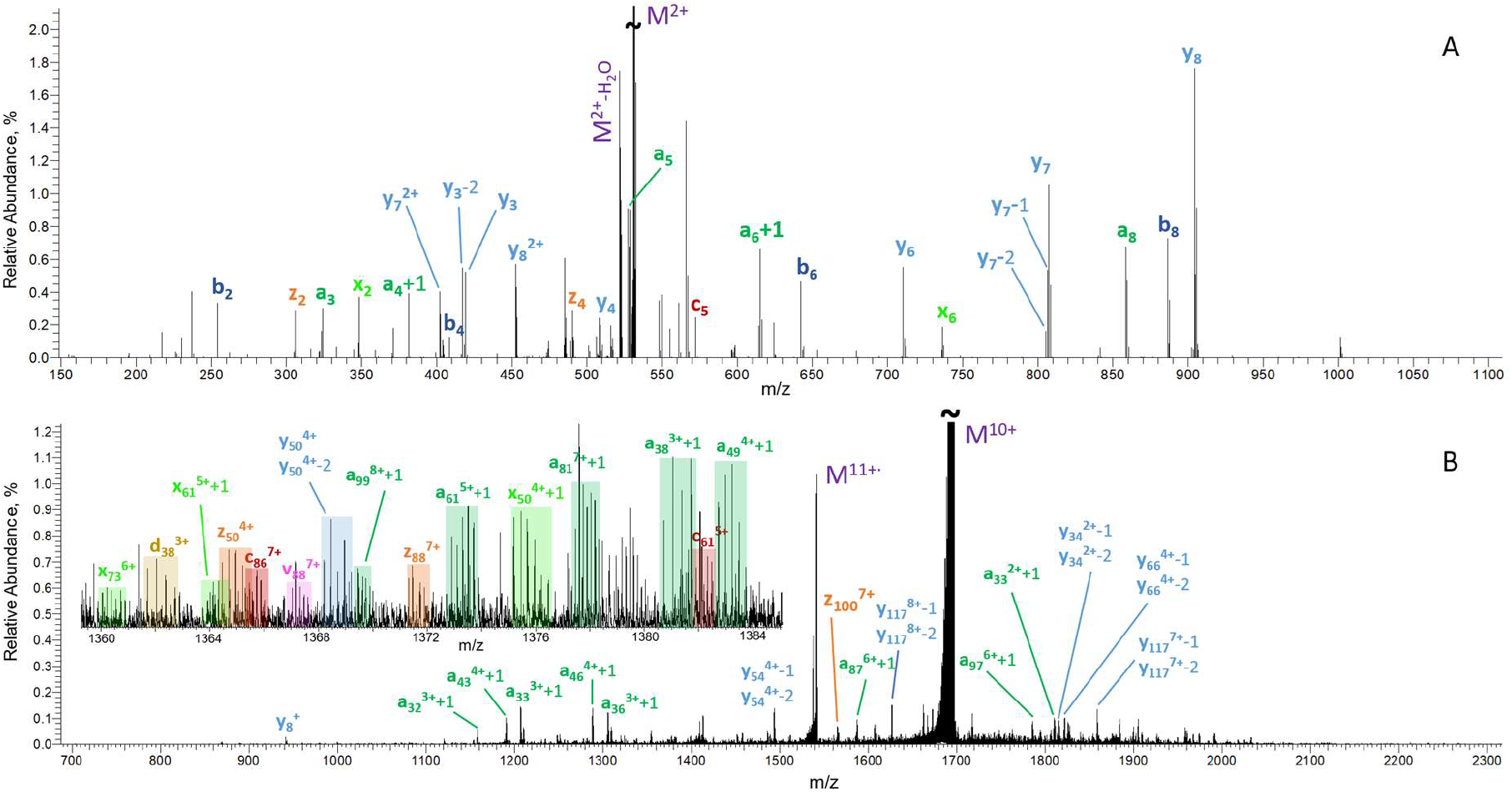
UVPD spectra of bradykinin^2+^ (A) and myoglobin^10+^(B), acquired following three pulses of the 193 nm UV light with the energy of 5 mJ per pulse. AGC values were set to one million for both analytes. For clarity, only few selected fragments are annotated in the spectrum of myoglobin^10+^(B). The inset in (B) contains the assignment of all fragments identified in the region between m/z 1360 and 1384.

Conversion of precursor to fragments with UVPD is generally suboptimal and we asked the question would supplemental activation with IRMPD improve fragment yields. A series of experiments was performed on myoglobin^10+^ and [carbonic anhydrase]^20+^ using IR-activation methods with a range of IR laser power outputs (Figures S15-S18). The mass-selected precursor was either preactivated by IR radiation prior to the UVPD, or coactivated continuously by IR radiation during triggering of three UV pulses. The analysis of resulting mass spectra demonstrated that both approaches had either no improvement or were detrimental to the sequence coverage of the protein and intensities of fragments. The only exceptions to this observation were the increased intensities of *y* fragments for carbonic anhydrase and *b, y*, and *c* fragments for myoglobin. The increased yields of *b* and *y* fragments are most likely due to IR-induced fragmentation of the precursor. The generation of *c* fragments is a radical-driven reaction and would be a product of UVPD where the supplementary IRMPD dissociates the high-order structure of the precursor. These results are similar to those obtained for IR-activated UVPD of ubiquitin.^59^

### Performance of ECD and EID

Omnitrap ECD performance has been previously described.^49^ We extended the application of ECD to incorporate IR supplemental activation which has previously been shown to be beneficial.^26-37^ Ions were transferred to segment Q5 and irradiated by low-energy (1-2 eV) electrons. Initial work involved ubiquitin^8+^, myoglobin^10+^, myoglobin^20+^, and [carbonic anhydrase]^20+^, and we found 50, 30, 30, and 20 ms of irradiation, respectively, was optimal and creates prominent charge-reduced species (Figure 4A, S19). We observed series of c/z and a+1/y fragments (Figures S20, S21A), with the latter pair resulting from the ECD-induced migration of H· to amide nitrogen, which is less favourable than association of H· with carbonyl carbon^10^. The sequence coverages were ranging from 97% and 89% for ubiquitin^8+^ and myoglobin^20+^ to 61% and 44% for [carbonic anhydrase]^20+^ and myoglobin^10+^, respectively (Figures S20, S21A), reflecting the dependence of the efficiency of ECD on charge density of precursor.^60^ To demonstrate the effect of IR-activation we chose myoglobin^10+^ as it has the lowest sequence coverage but prominent charge-reduced precursor peaks in ECD alone. The preactivation by IRMPD of myoglobin^10+^ prior to ECD in the Omnitrap leads to significantly higher sequence coverage by a/c/z fragments (Figures 4B, S21B, S22).

**Figure 4.**
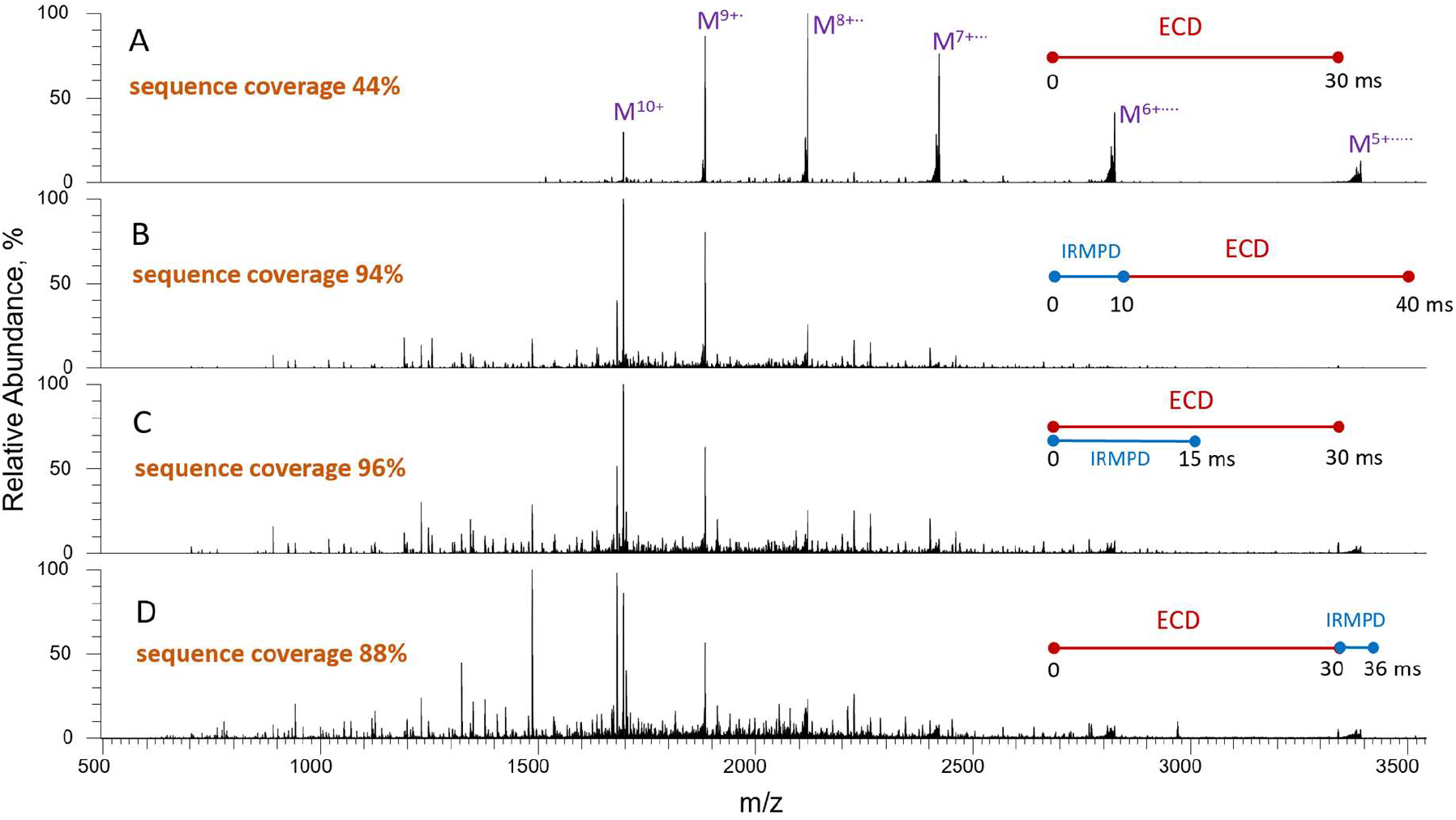
ECD and IR-activated-ECD spectra of myoglobin^10+^. A) ECD 30 ms; B) IRMPD 10 ms (23 % laser duty cycle) followed by 30 ms ECD; C) IRMPD (14 % laser duty cycle) concurrently with ECD for 15 ms followed by 15 ms of ECD; D) ECD 30 ms followed by 6 ms IRMPD (23 % laser duty cycle).

**Figure 5.**
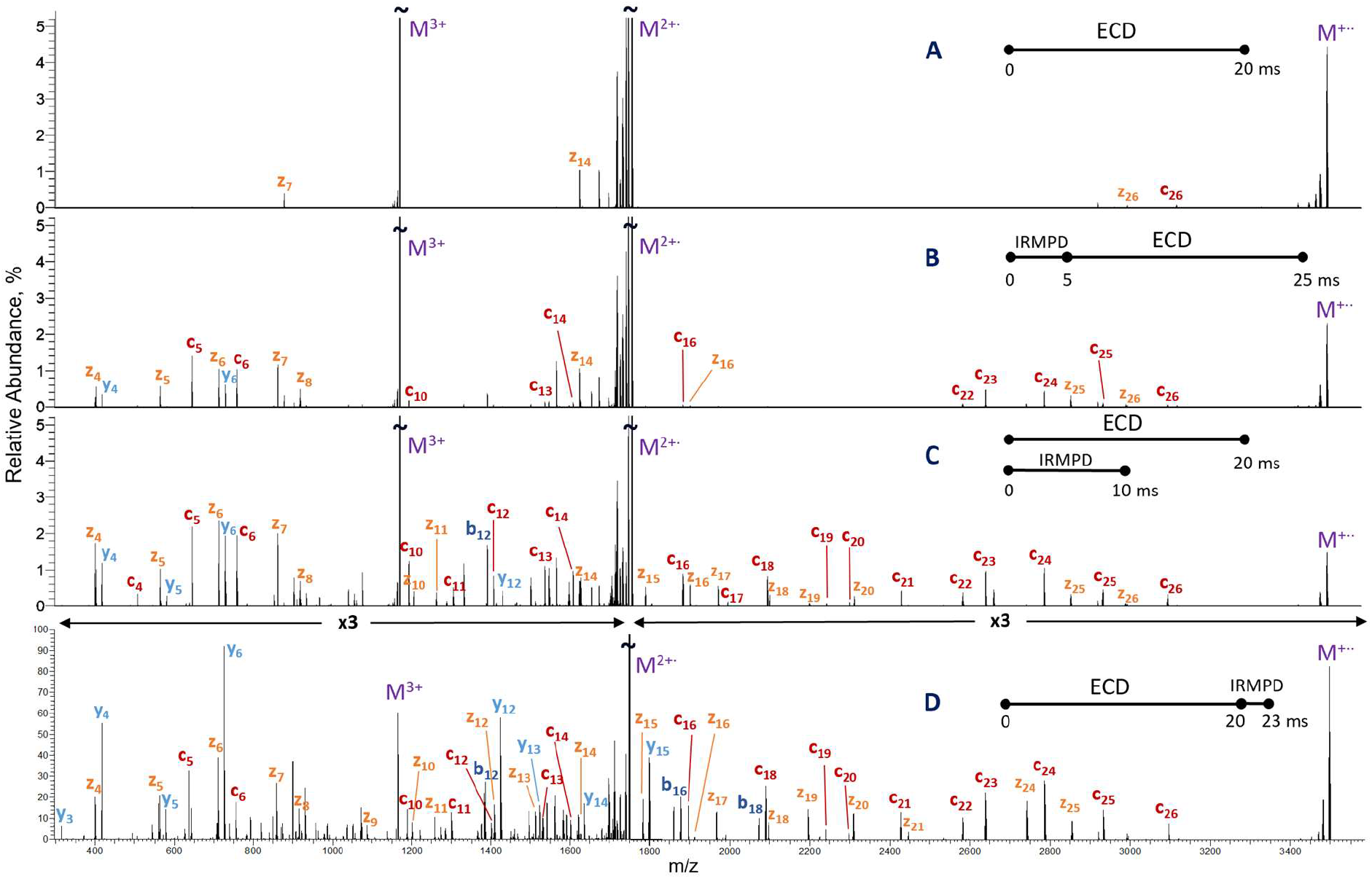
ECD and IR-activated-ECD spectra of triply charged chain B of insulin. A) ECD 20 ms; B) IRMPD 5 ms (21 % laser duty cycle) followed by 20 ms; C) simultaneous irradiation of precursor ions by IR (15 % laser duty cycle) and ECD for 10 ms followed by 10 ms of ECD; D) ECD 20 ms followed by 3 ms IRMPD (26 % laser duty cycle)

The number of identified fragments and total sequence coverage further increase when the precursor is irradiated by IR light and electrons simultaneously (Figure 4C, S21C, S22). In this experiment, the precursor was coactivated by IR only during the first half of the ECD reaction, because longer exposure to IR radiation led to extensive primary and secondary fragmentation and loss of signal even with the minimal power output of the laser. When coactivated using just 14% of the laser duty cycle, ECD yielded 90% and 72% of the sequence covered by *c* and *z* fragments respectively, which summed to 96% of the sequence coverage in total (Figures S21C, S22). This result resembles that obtained previously in AI-ETD of myoglobin in a modified Orbitrap HCD collision cell^35^ and linear ion trap^36^. The postactivation of ECD fragments by IR light leads to higher numbers of identified *c* and *z* fragments and higher sequence coverage compared to ECD alone but loses out to pre- and co-activation methods (Figures S21D, S22). The increased yield of *c* and *z* fragments in this latter approach can be attributed to the disruption of charge-reduced complexes by IR; however, the secondary activation by IR dissociates primary fragments as well thus reducing the total efficiency of IR-postactivation of ECD. As a whole, all three activation methods increased the numbers of *c* and *z* fragments compared to ECD alone, with preactivation being the most efficient for the yield of *z* fragments, and coactivation favouring the generation of *c* fragments (Figure S22).

We analysed the usefulness of electron ionisation dissociation (EID) and activated EID on the same low-charge-state precursor myoglobin^10+^. As shown previously, Omnitrap-EID provides a complete sequence coverage for proteins as small as ubiquitin^49^. In our experiments, irradiation of myoglobin^10+^ for 30 ms by 35 eV electrons resulted in 82% sequence coverage (Figures S23-S25). The pre- and co-activation by low-energy IR increased the yields of all ions of main series, which resulted in slightly better sequence coverage of 91% and 93%, respectively. The postactivation by IR leads to the reduced numbers of identified fragments of all types compared to EID alone probably due to secondary fragmentation, with the exception of increased numbers of IR-induced *b* and *y* ions.

The IR-activated ECD of peptides followed the similar trends as that of proteins. We found 20 ms was optimal irradiation time for peptides, and the charge-reduced species were the main products in ECD of [Glu-fibrinopeptide B]^2+^ and triply charged chain B of insulin (Figures S26A, 5A). The activation of precursor with low-power IR prior to ECD moderately increased the number of identified fragments of *c*/*z* type (Figures S26B, 5B). Co-activation with IRMPD further increased the numbers and intensities of *c* and *z* ions and led to the appearance of additional *b* and *y* fragments (Figures S26C, 5C). The increased yields of *c, z, b*, and *y* fragments was also observed when precursor and ECD products were postactivated by low-energy IR (Figures S27, S28). Overall, similar to IR-activated ECD of proteins, the number of cleaved bonds and intensities of *c* and *z* fragments of peptides dramatically increased when the precursors were pre-, co-, or post-activated by IR. Among these three activation methods, the coactivation by IR yields highest *c, z, b*, and *y* ion currents (Figure S28), but intensities of each individual *c* or *z* fragment can reach their maximum values in either co-, or post-activation by IR (Figure S27).

The main goal of pre-, co-, and post-activation of ECD by low-power IR is to increase the yields of *c* and *z* fragments. As a result, a significant part of precursor with low charge state typically remains unfragmented. This precursor can be further dissociated, for example, in CID or HCD to generate intensive complementary *b* and *y* fragments, that increase the sequence coverage and confidence of identification of a peptide or protein; this approach has been implemented in EThcD.^61, 62^ In a similar way, the use of high-power IRMPD after ECD consumed the majority of the remaining precursor and produced near-complete series of *z* and *y* ([Glu-fibrinopeptide B]^2+^) or *c, z*, and *y* (triply charged insulin chain B) fragments (Figures S26D, 5D).

The total length of a single ECD or coactivated ECD experiment in the Omnitrap amounts to 40 ms + N, where N is irradiation time in ms. Thus, in ECD experiments of peptides, the length of a single scan amounts to 60 ms not including transfer time within the Exploris instrument, which is few times shorter than scan lengths reported for ECD implemented in electromagnetostatic cell^15^ and digital quadrupole ion trap^19^ and similar to ECD pulse lengths used in ICR Penning trap^63^. Such relatively short scan time makes Omnitrap-ECD suitable for analysis of PTMs in complex peptide mixtures on the LCMS scale.

## Conclusion

The efficiency of UVPD in the Omnitrap is comparable with results reported in literature, as exemplified by sequence coverages of proteins with molecular weights <30 kDa. High-energy process UVPD doesn’t benefit from IR-activation, suggesting that energising vibrational modes does not affect the dissociation pathways in this type of fragmentation. Low-energy (1-2 eV) electrons were used for efficient ECD of peptides and proteins. Pre-, co-, and post-activation of a low charge state precursors by IR light leads to significant increase sequence coverage, as in the case of myoglobin^10+^, for which near-complete sequence coverage was obtained. We demonstrated the ability of the Omnitrap platform coupled to an Orbitrap mass spectrometer to perform efficient UVPD and ECD and their combination with IRMPD for analysis of peptides and top-down analysis of proteins.

## Associated content

### Supporting Information

The supporting information is available free of charge at the <u>ACS website</u>

Schematic drawing of the instrument and propagating IR laser beam (Figure S1); schematic drawing of the overlap between ion cloud and IR laser beam in Q5 (Figure S2) or UV laser beam in Q8 (Figure S3); pressure profile in the Omnitrap (Figure S4); the dependence of the efficiency of IRMPD, trapping efficiency, and depletion of precursor on delay after gas pulse (Figure S5), *q* value (Figure S6), and irradiation time (Figure S7); IRMPD of [Glu-fibrinopeptide B]^2+^ in Q2, Q5, and Q8 (Figure S8); IRMPD spectra of proteins (Figure S9); dependence of UVPD efficiency on the number of UV laser pulses for bradykinin^2+^ (Figure S10) and ubiquitin^8+^ (Figure S11); UVPD spectra of ubiquitin^8+^ and [carbonic anhydrase]^20+^ (Figure S12); distribution of numbers of main-series fragments identified in UVPD of ubiquitin^8+^ (Figure S13); sequence coverage maps for UVPD of proteins (Figure S14); spectra of IR-activated UVPD of myoglobin^10+^ (Figure S15) and [carbonic anhydrase]^20+^ (Figure S17); ion currents of various types of fragments in IR-activated UVPD of myoglobin^10+^ (Figure S16) and [carbonic anhydrase]^20+^ (Figure S18); spectra (Figure S19) and sequence coverage maps (Figure S20) for ECD of proteins; sequence coverage maps (Figure S21) and sequence coverage distributions (Figure S22) for IR-activated ECD of myoglobin^10+^; spectra (Figure S23) and corresponding sequence coverage maps (Figure S24) and sequence coverage distributions (Figure S25) for IR-activated EID of myoglobin^10+^; spectra of IR-activated ECD of [Glu-fibrinopeptide B]^2+^ (Figure S26); ion currents of fragments for IR-activated ECD of [Glu-fibrinopeptide B]^2+^ (Figure S27) and triply charged chain B of insulin (Figure S28); details about analytes used in the experiments (Table S1); types of fragments searched in the spectra using the in-house software and MS-TAFI (Table S2) (PDF)

## Author Information

## Author Contributions

A.S., K.F., A.M., D.P. and S.M. designed the instrument. A.S. and D.M. built the Omnitrap A.S., K.F., A.M., D.P. and S.M. commissioned the instrument. N.L. collected the data. N.L., M.K., D.P. and S.M. analysed the data. N.L. and S.M. wrote the initial draft. All authors have revised the manuscript and have given approval to its final version.

‡These authors contributed equally

## Notes

The authors declare no competing financial interest.

## Acknowledgements

The Next Generation Chemistry theme at the Franklin Institute is supported by the EPSRC (V011359/1 (P)).

## For Table of Contents Only

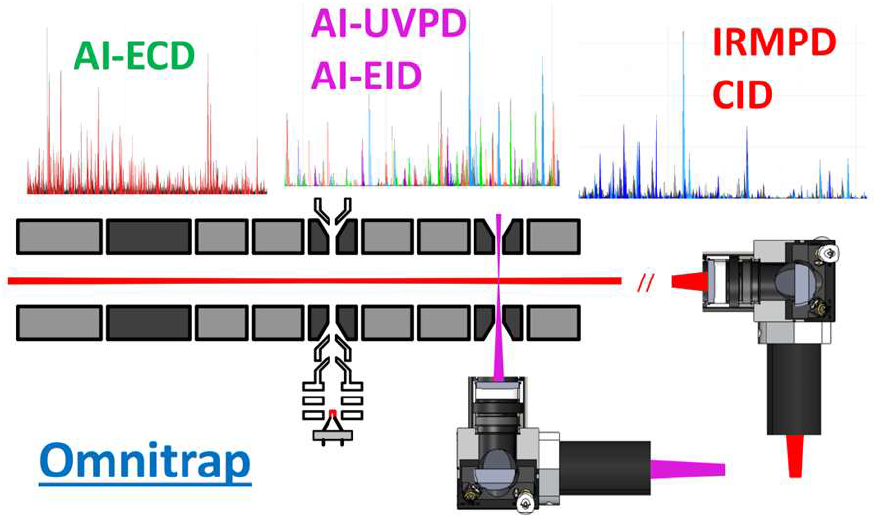

## Notes

### Competing Interest Statement

The authors have declared no competing interest.

