## Supplementary Information for "Characterisation of an Omnitrap-Orbitrap platform equipped with IRMPD, UVPD and ExD for the analysis of peptides and proteins"

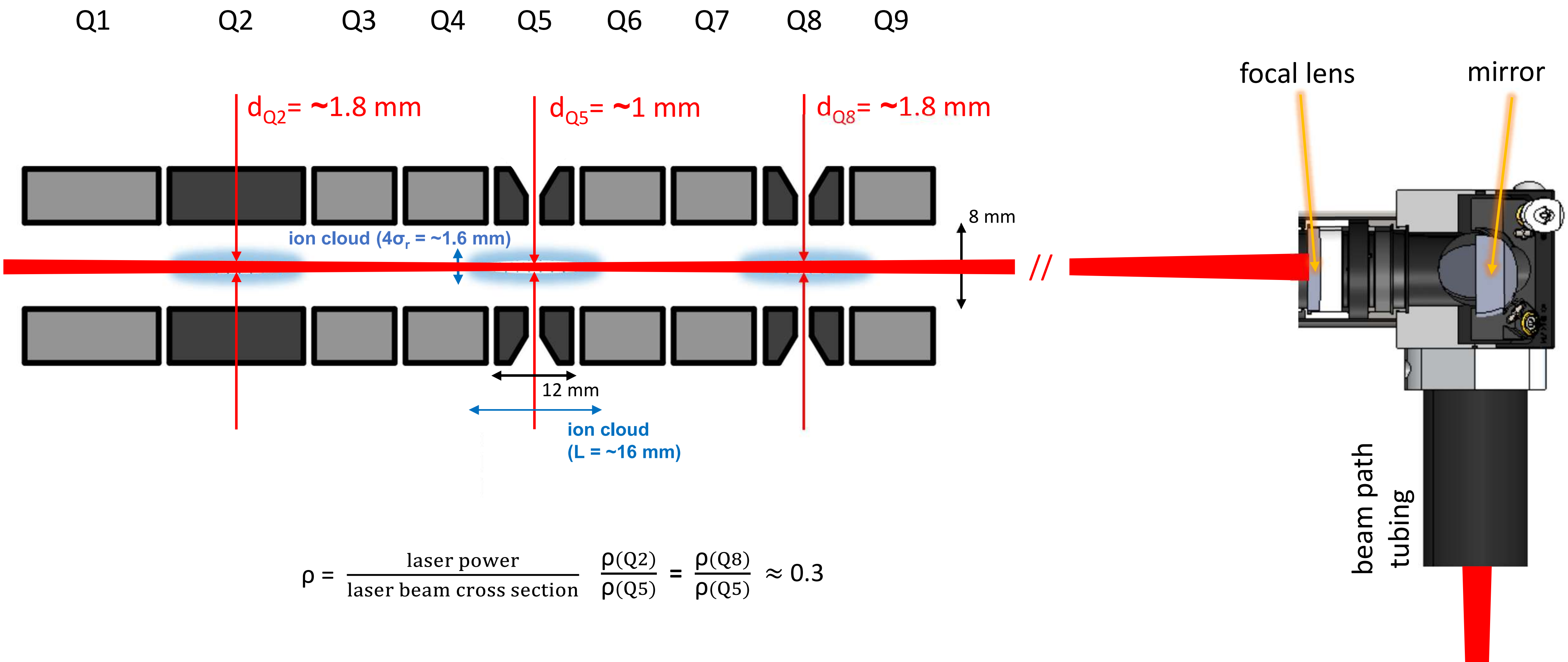


Figure S1. Schematic drawing of the cross-section of the Omnitrap in the plane parallel to its axis and the overlap between an ion cloud (500000 charges) trapped at q=0.15 in Q2/Q5/Q8 and an IR laser beam. The radial density of the ion cloud is represented by a normal distribution with a variance (σ) of ~0.4 mm. The variation corresponding to 4σ covers approximately 99% of the population of charges. The ideal case of a laser beam perfectly aligned along the axis is considered.

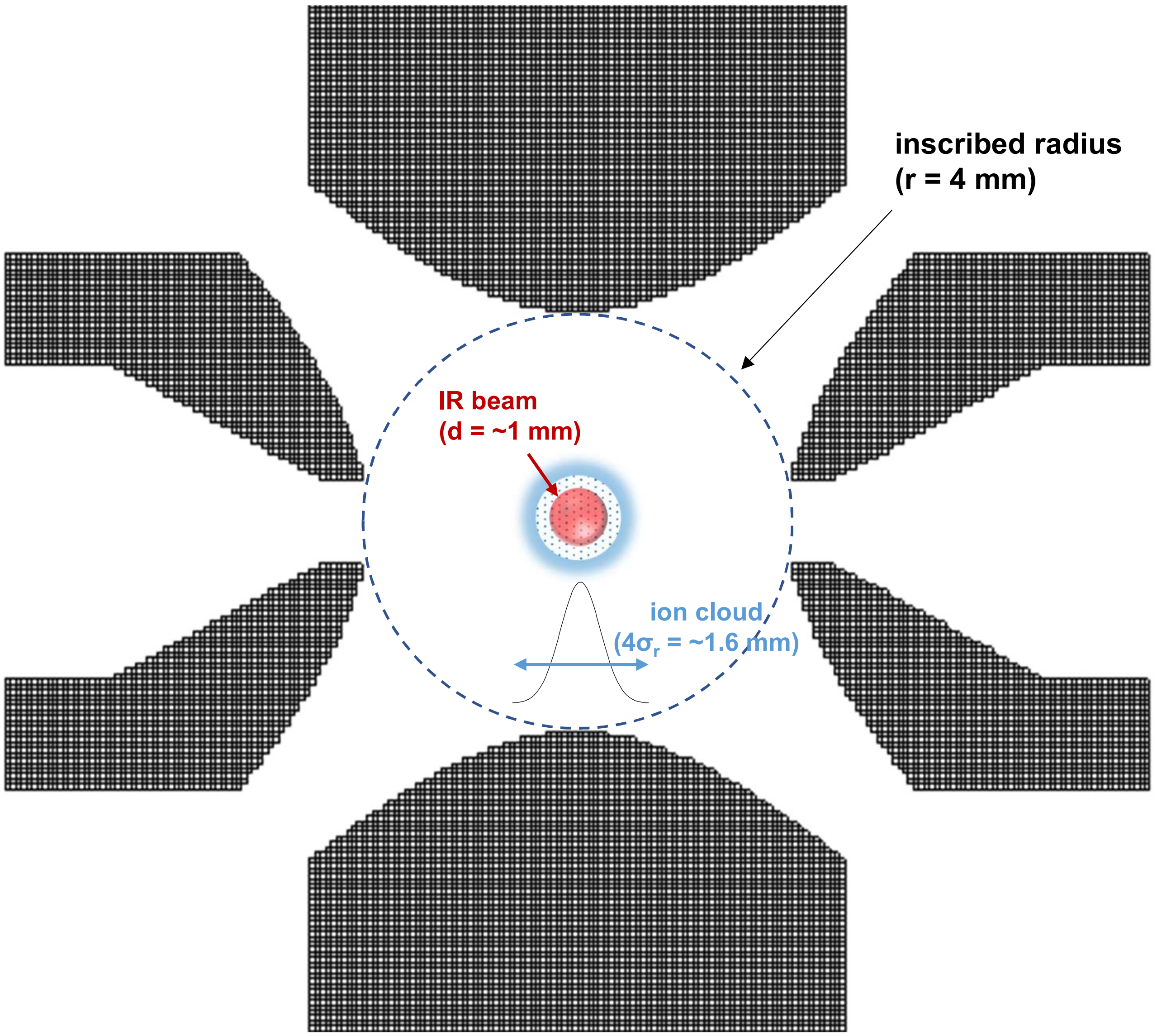


Figure S2. Schematic drawing of the cross-section of the segment Q5 in the plane orthogonal to the axis of the trap and the overlap between an ion cloud (500000 charges) trapped at q=0.15 and an IR laser beam. The radial density of the ion cloud is represented by a normal distribution with a variance (σ) of ~0.4 mm. The variation corresponding to 4σ covers approximately 99% of the population of charges.

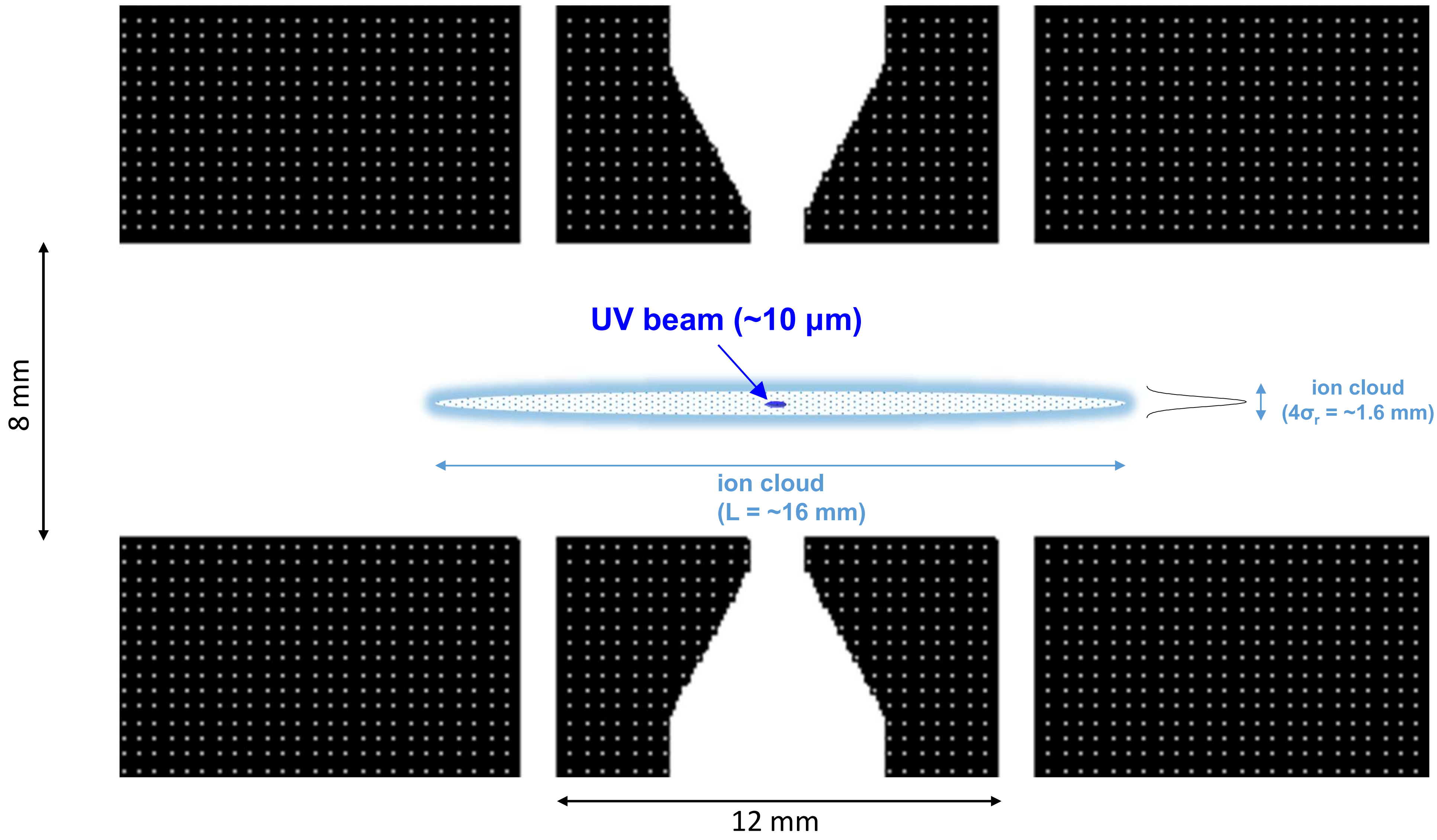


Figure S3. Schematic drawing of the cross-section of the segments Q7, Q8, Q9 in the plane parallel to the axis of the trap and the overlap between an ion cloud (500000 charges) trapped at q=0.15 and a UV laser beam. The radial density of the ion cloud is represented by a normal distribution with a variance (σ) of ~0.4 mm. The variation corresponding to 4σ covers approximately 99% of the population of charges.

Table S1. Details about analytes used in the experiments

| analyte | sequence | modification | concentration/solvent | monoisotopic mass |
|---|---|---|---|---|
| bradykinin | RPPGFSPFR | - | 1 µM in 50:25:24:1 v/v ACN:MeOH:$H_2O$:acetic acid | 1059.56 |
| Glu-fibrinopeptide B | EGVNDNEEGFFSAR | - | 0.1 µM in 49.5:49.5:1 ACN:water:formic acid | 1569.67 |
| insulin chain B | FVNQHLCGSHLVEALYLVCGERGFFYTPKA | Trioxidation of Cys residues | 1 µM in 49.5:49.5:1 ACN:water:formic acid | 3493.64 |
| ubiquitin | MQIFVKTLTGKTITLEVEPSDTIENVKAKIQD KEGIPPDQQRLIFAGKQLEDGRTLSDYNIQKE STLHLVLRLRGG | - | 0.5 µM in 49.5:49.5:1 ACN:water:acetic acid | 8559.62 |
| myoglobin | GLSDGEWQQVLNVWGKVEADIAGHGQEVLI RLFTGHPETLEKFDKFKHLKTEAEMKASEDL KKHGTVVLTALGGILKKKGHHEAELKPLAQS HATKHKIPIKYLEFISDAIIHVLHSKHPGDFGA DAQGAMTKALELFRNDIAAKYKELGFQG | - | 0.1 µM in 49.5:49.5:1 ACN:water:formic acid | 16983.98 |
| carbonic anhydrase | SHHWGYGKHNGPEHWHKDFPIANGERQSPV DIDTKAVVQDPALKPLALVYGEATSRRMVNN GHSFNVEYDDSQDKAVLKDGPLTGTYRLVQF HFHWGSSDDQGSEHTVDRKKYAAELHLVHW NTKYGDFGTAAQQPDGLAVVGVFLKVGDANP ALQKVLDALDSIKTKGKSTDFPNFDPGSLLPN VLDYWTYPGSLTTPPLLESVTWIVLKEPISVS SQQMLKFRTLNFNAEGEPELLMLANWRPAQ PLKNRQVRGFPK | N-term acetylation | 10 µM in 49.5:49.5:1 ACN:water:formic acid | 29006.68 |

Table S2. Types of fragments searched in the spectra using the in-house software and MS-TAFI

| **fragmentation technique** | **types of fragments searched (in-house software)** | **types of fragments searched (MS-TAFI)** |
|---|---|---|
| IRMPD | b, y | b, y (CID option) |
| UVPD<br>IR-activated-UVPD | a, a+1, b, b+1, b+2, c, x, x+1, y, y-1, y-2, z | a, a+1, b, c, x, x+1, y, y-1, y-2, z (UVPD option) |
| ECD | a, a+1, c, c+1, c-1, y, z, z+1, z-1 | a, a+1, b, c, x, x+1, y, y-1, y-2, z (UVPD option) |
| IR-activated-ECD | a, a+1, b, c, c+1, c-1, y, z, z+1, z-1 | a, a+1, b, c, x, x+1, y, y-1, y-2, z (UVPD option) |
| EID<br>IR-activated-EID | a, a+1, b, c, c-1, c+1, x, x+1, y, z, z-1, z+1 | a, a+1, b, c, x, x+1, y, y-1, y-2, z (UVPD option) |

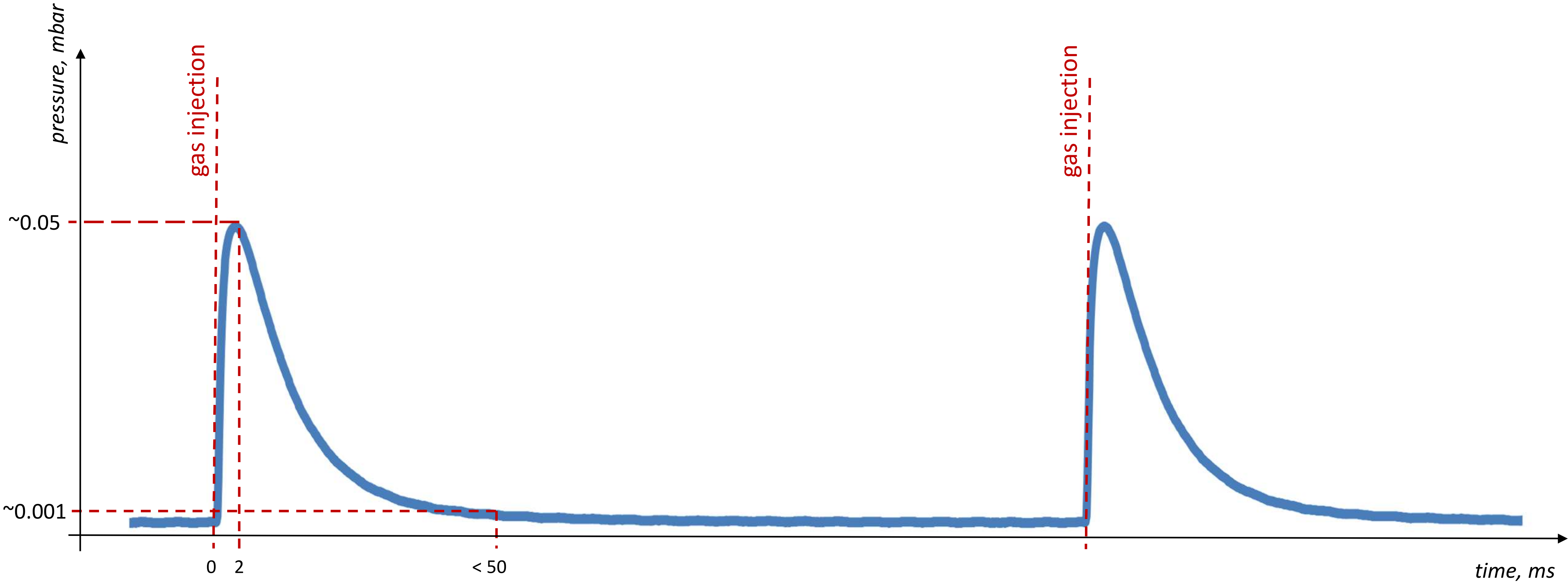


Fig S4. Pressure profile in the Omnitrap following a pulse of buffer gas. The temporal profile of pressure is controlled by adjusting the width of the voltage pulse applied to the solenoid valve. A maximum pressure of 0.05 mbar is reached within 2 ms from releasing nitrogen gas into the ion trapping region, while the period where pressure is sufficiently high to promote collisional cooling (>0.001 mbar) can be extended up to ~50 ms.

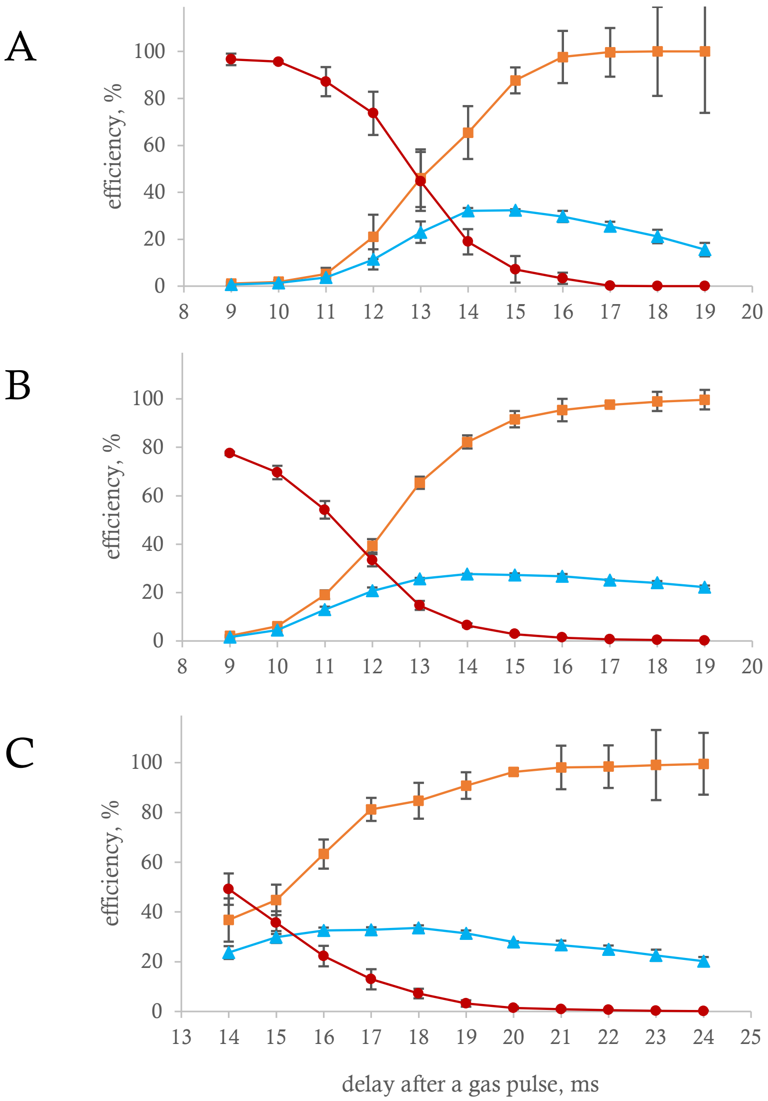


| | Q2 | Q5 | Q8 |
|---|---|---|---|
| irrad. time, ms | 11 | 4 | 12 |
| *q* value | 0.2 | 0.2 | 0.2 |
| laser power, % duty cycle | 30 | 30 | 30 |

$$\text{photodiss. eff.} = \frac{F}{F + P}$$

$$\text{ion trapping eff.} = \frac{F}{P_0}$$

$$\text{norm. int. of precursor} = \frac{P}{P_0}$$

F – summed intensities of all fragments
P – remaining intensity of precursor after fragmentation
$P_0$ – intensity of precursor before fragmentation

Fig S5. The dependence of the photodissociation efficiency, trapping efficiency and normalised intensity of the precursor on the delay between a gas pulse and IR triggering in the segments Q2 (A), Q5 (B), and Q8 (C) in the IRMPD of [Glu-fibrinopeptide B]$^{2+}$. The parameters of the experiments are given in the table next to the charts. Each data point represents an average of a triplicate with standard deviations given as error bars.

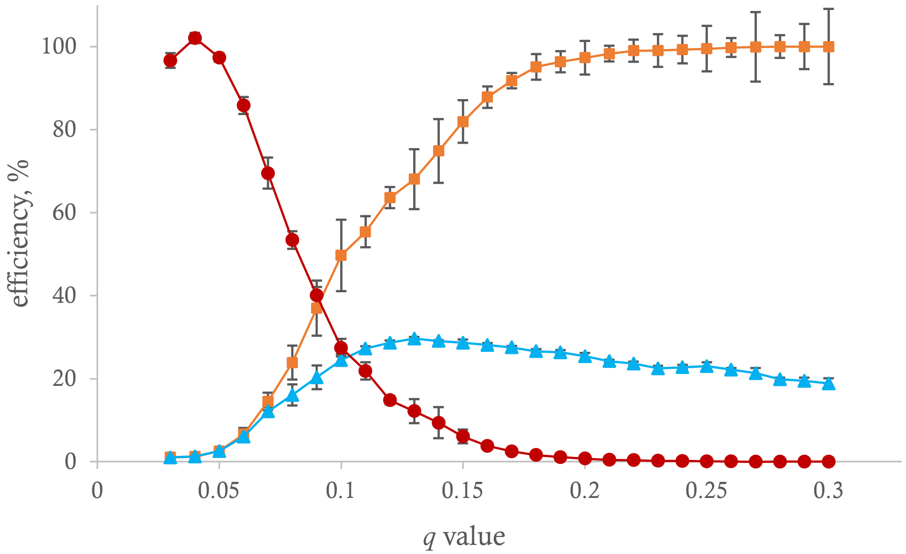


| | Q5 |
|---|---|
| irrad. time, ms | 5 |
| delay after a gas pulse | 13 |
| laser power, % duty cycle | 30 |

$$\text{photodiss. eff.} = \frac{F}{F + P}$$

$$\text{ion trapping eff.} = \frac{F}{P_0}$$

$$\text{norm. int. of precursor} = \frac{P}{P_0}$$

F – summed intensities of all fragments
P – remaining intensity of precursor after fragmentation
$P_0$ – intensity of precursor before fragmentation

Fig S6. The dependence of the photodissociation efficiency, trapping efficiency and normalised intensity of the precursor on the *q* value of the Omnitrap in the segment Q5 in the IRMPD of [Glu-fibrinopeptide B]$^{2+}$. The parameters of the experiment are given in the table next to the chart. Each data point represents an average of a triplicate with standard deviations given as error bars.

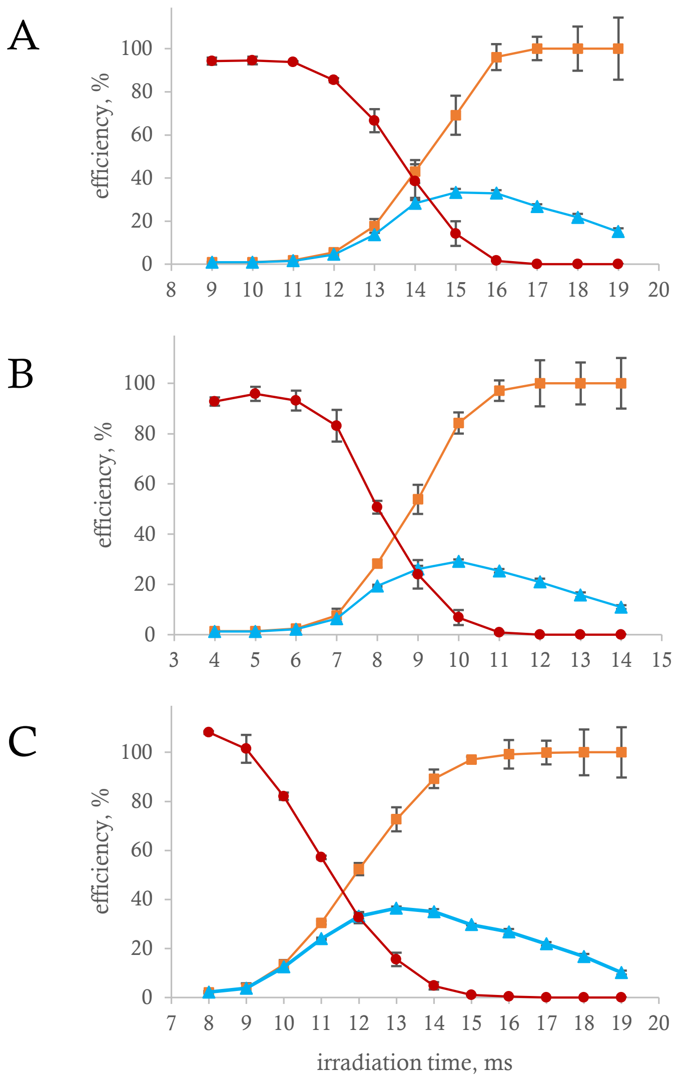


■ photodissociation efficiency
▲ ion trapping efficiency
● normalised intensity of the precursor

| | Q2 | Q5 | Q8 |
|---|---|---|---|
| laser power, % duty cycle | 30 | 20 | 30 |
| *q* value | 0.2 | 0.2 | 0.2 |
| delay after a gas pulse | 10 | 10 | 15 |

$$\text{photodiss. eff.} = \frac{F}{F + P}$$

$$\text{ion trapping eff.} = \frac{F}{P_0}$$

$$\text{norm. int. of precursor} = \frac{P}{P_0}$$

F – summed intensities of all fragments
P – remaining intensity of precursor after fragmentation
$P_0$ – intensity of precursor before fragmentation

Fig S7. The dependence of the photodissociation efficiency, trapping efficiency and normalised intensity of the precursor on the length of IR irradiation in the segments Q2 (A), Q5 (B), and Q8 (C) in the IRMPD of [Glu-fibrinopeptide B]$^{2+}$. The parameters of the experiments are given in the table next to the charts. Each data point represents an average of a triplicate with standard deviations given as error bars.

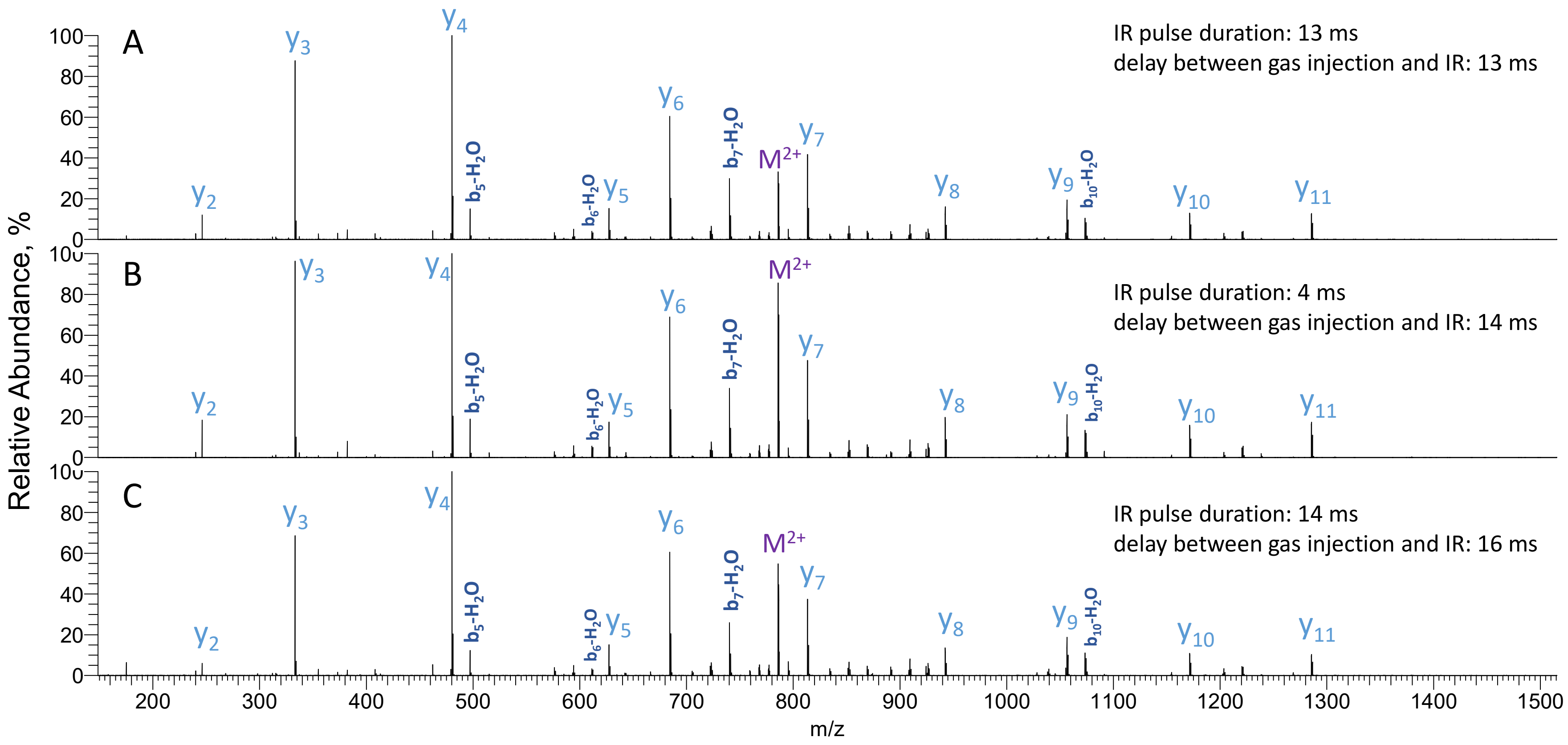


Fig S8. IRMPD spectra of [Glu-fibrinopeptide B]$^{2+}$ acquired in the segments Q2 (A), Q5 (B) and Q8 (C). IRMPD spectra were acquired at q=0.2, 30% laser duty cycle. Laser pulse durations and delays between gas injection and IR pulsing are given in the figure

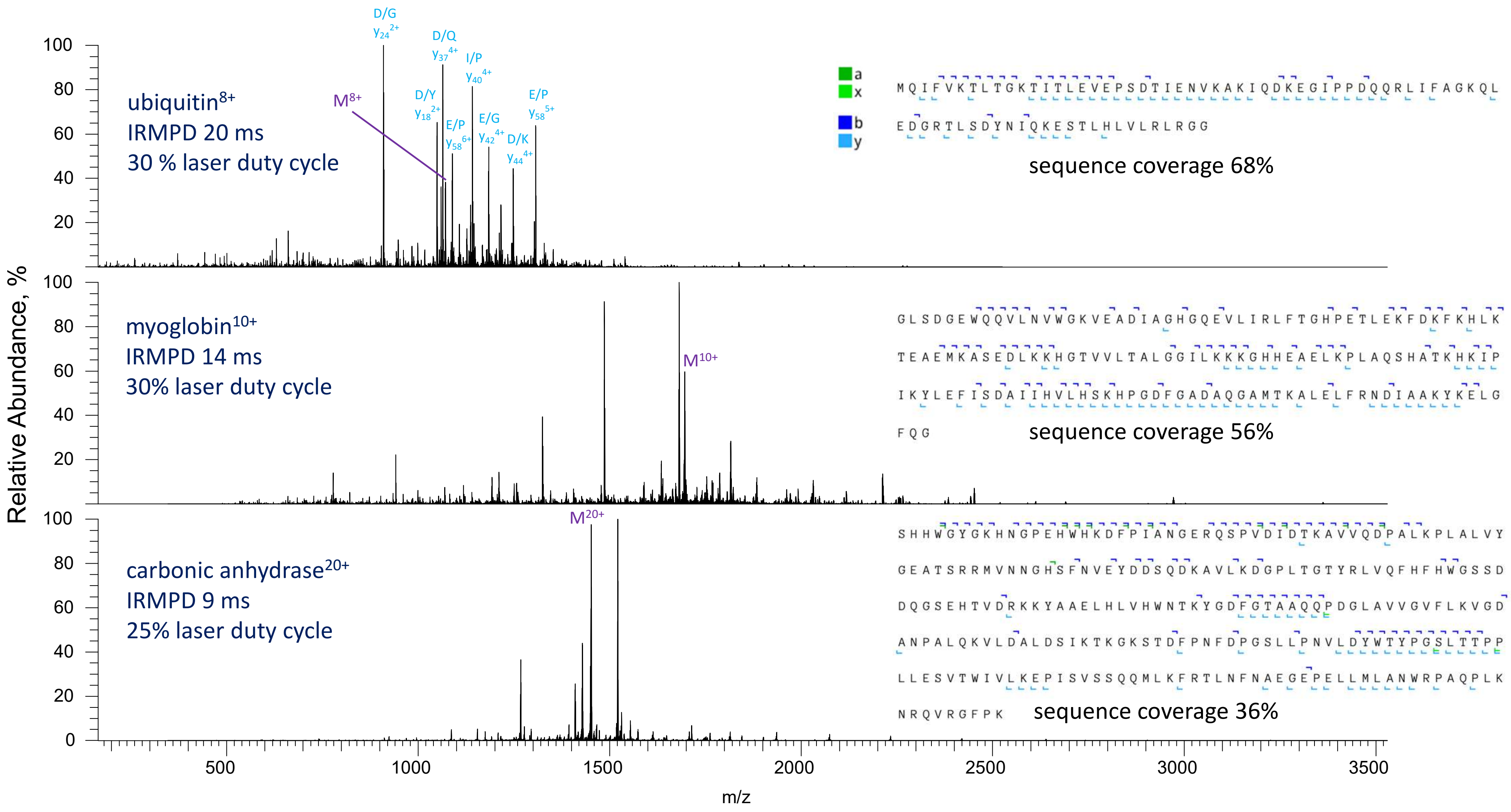


Fig S9. IRMPD spectra of ubiquitin$^{8+}$, myoglobin$^{10+}$ and [carbonic anhydrase]$^{20+}$ acquired in the segment Q5 of the Omnitrap at q=0.2 and corresponding fragment mapping. IRMPD irradiation time and laser power are given in the figure; the IR was triggered 11 ms after the gas pulse.

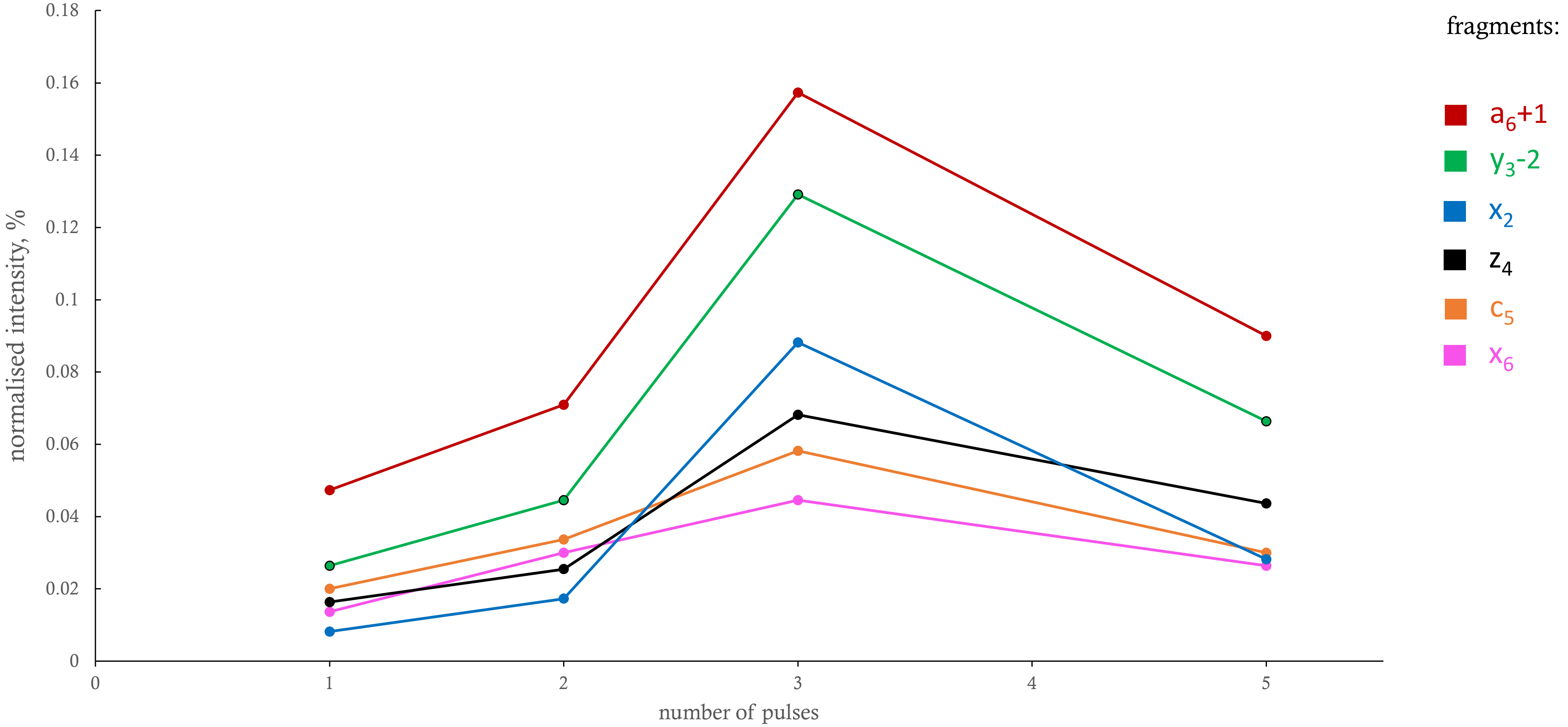


Fig S10. Intensities of few selected fragments normalised to the MS1 intensity of the precursor in the UVPD of $[\text{bradykinin}]^{2+}$ for different number of pulses of the UV laser. The energy of each pulse was 5 mJ.

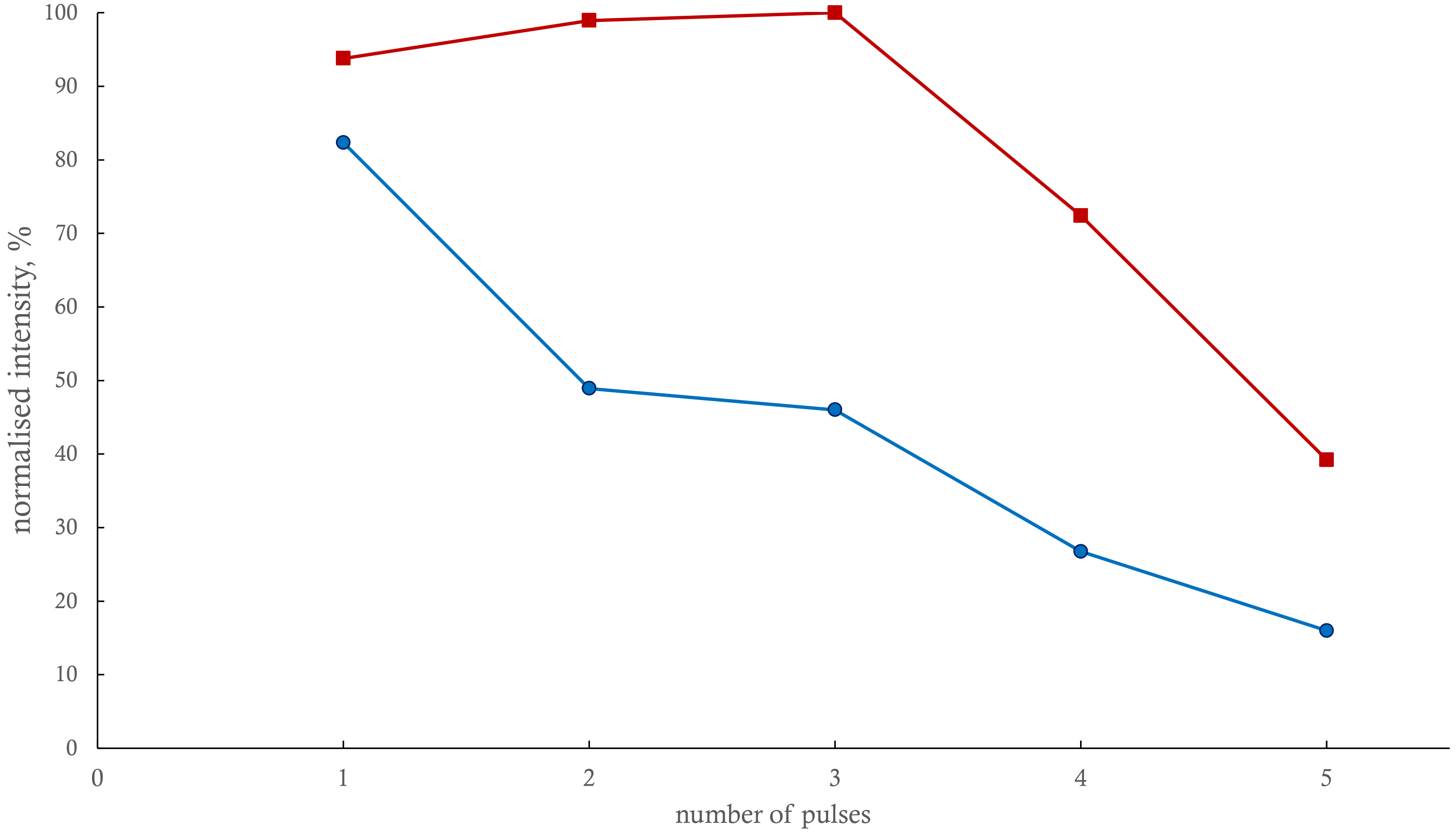


Fig S11. Total ion current of all main-series fragments of ubiquitin$^{8+}$ normalised to the highest value in the series (red) and intensity of unfragmented ubiquitin$^{8+}$ precursor normalised to its intensity in MS1 (blue) in the UVPD experiments for different number of pulses of the UV laser. The energy of each pulse was 5 mJ.

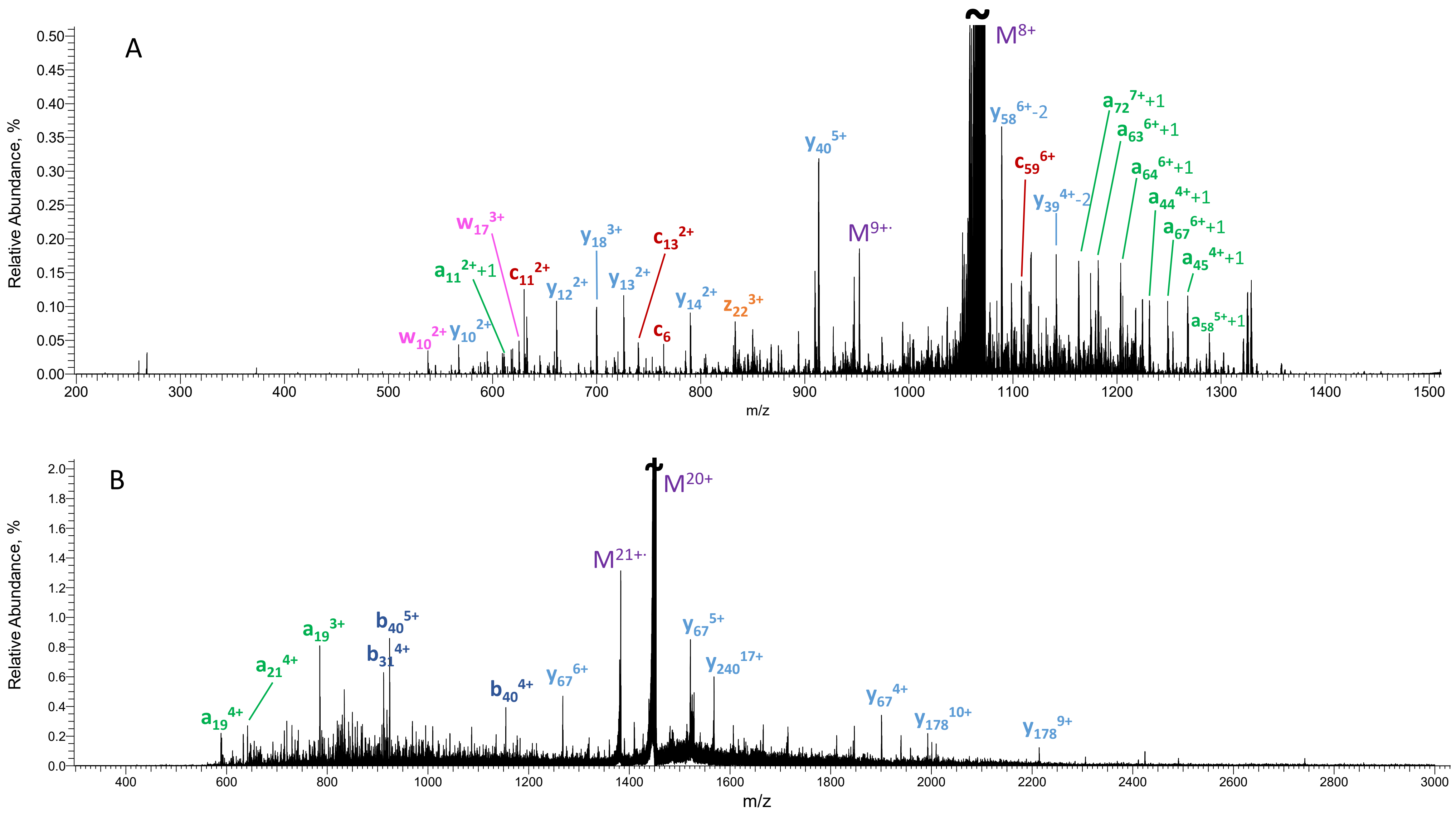


Fig S12. UVPD spectra of ubiquitin$^{8+}$ (A) and [carbonic anhydrase]$^{20+}$ (B) acquired following three pulses of the 193 nm UV laser with the energy of 5 mJ per pulse. For clarity, only few selected fragments are annotated

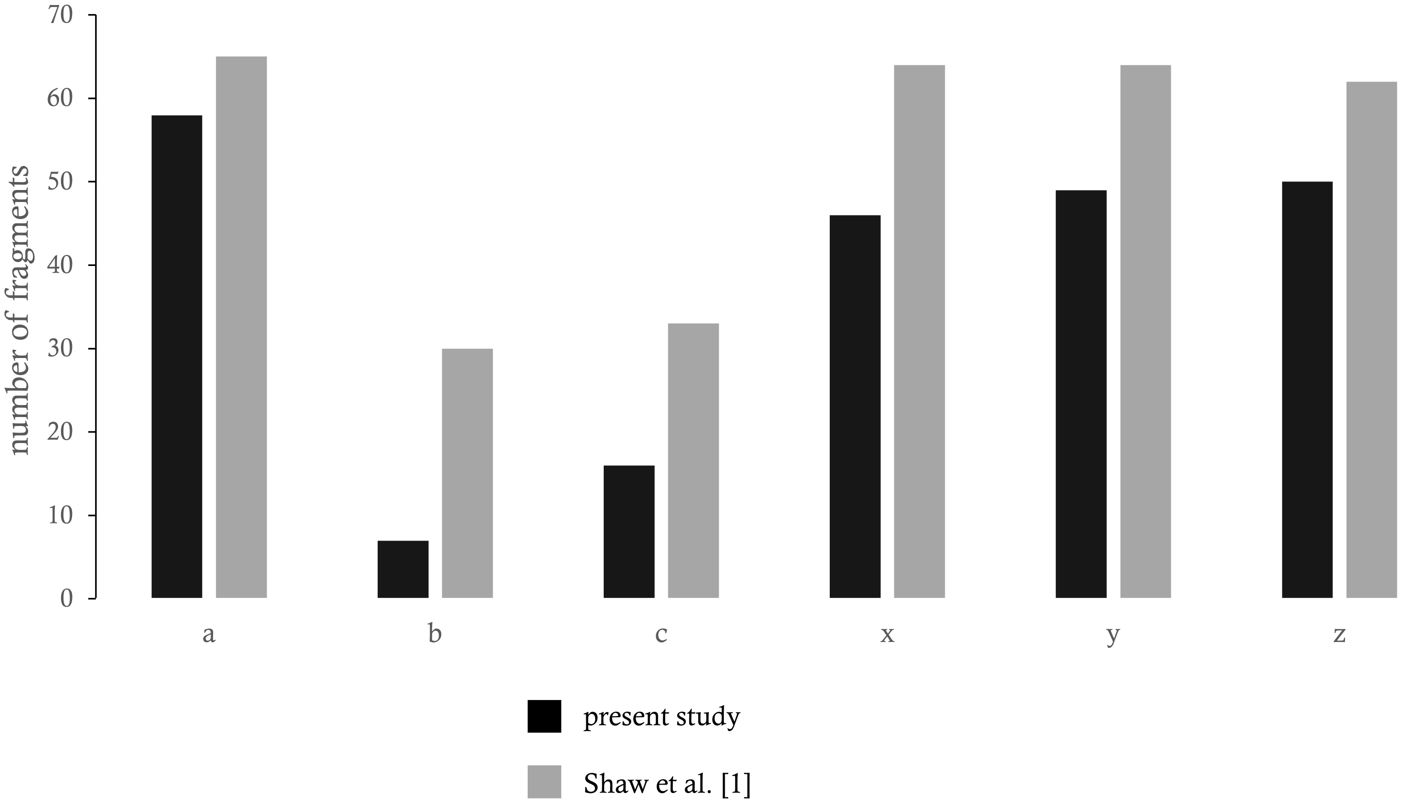


Fig S13. Number of fragments of different types identified in the UVPD experiments of ubiquitin$^{8+}$ reported in this study (3 pulses, 5 mJ/pulse, black) and [Ref 1] (1 pulse, 1-4 mJ/pulse, grey)

[1] Shaw, J. B.; Li, W.; Holden, D. D.; Zhang, Y.; Griep-Raming, J.; Fellers, R. T.; Early, B. P.; Thomas, P. M.; Kelleher, N. L.; Brodbelt, J. S. Complete protein characterization using top-down mass spectrometry and ultraviolet photodissociation. *Journal of the American Chemical Society* **2013**, *135* (34), 12646-12651.

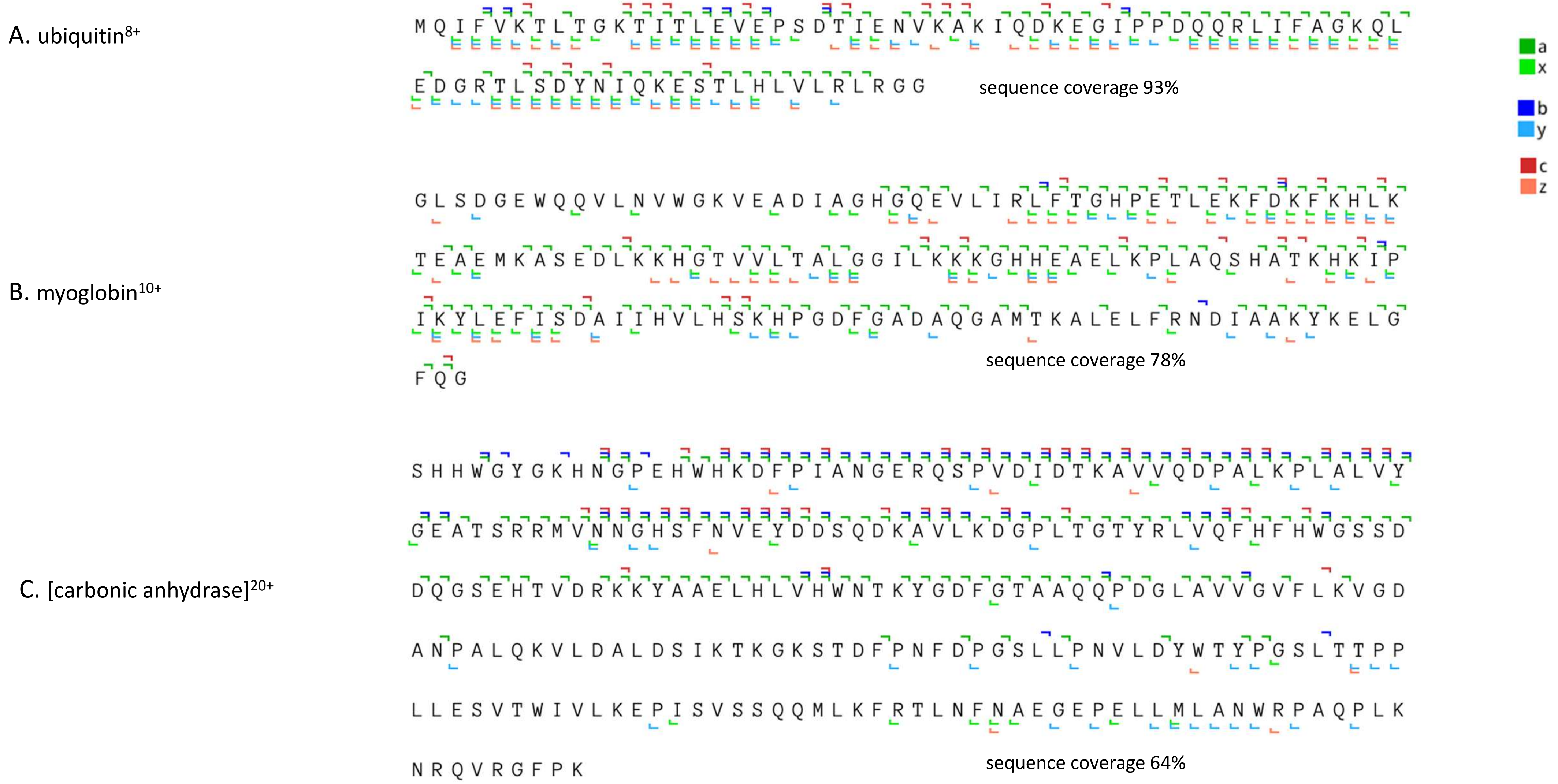


Fig S14. Fragment maps for UVPD experiments on ubiquitin$^{8+}$ (A), myoglobin$^{10+}$ (B), and [carbonic anhydrase]$^{20+}$ (C). The corresponding spectra were acquired following three pulses of the 193 nm UV laser with the energy of 5 mJ per pulse.

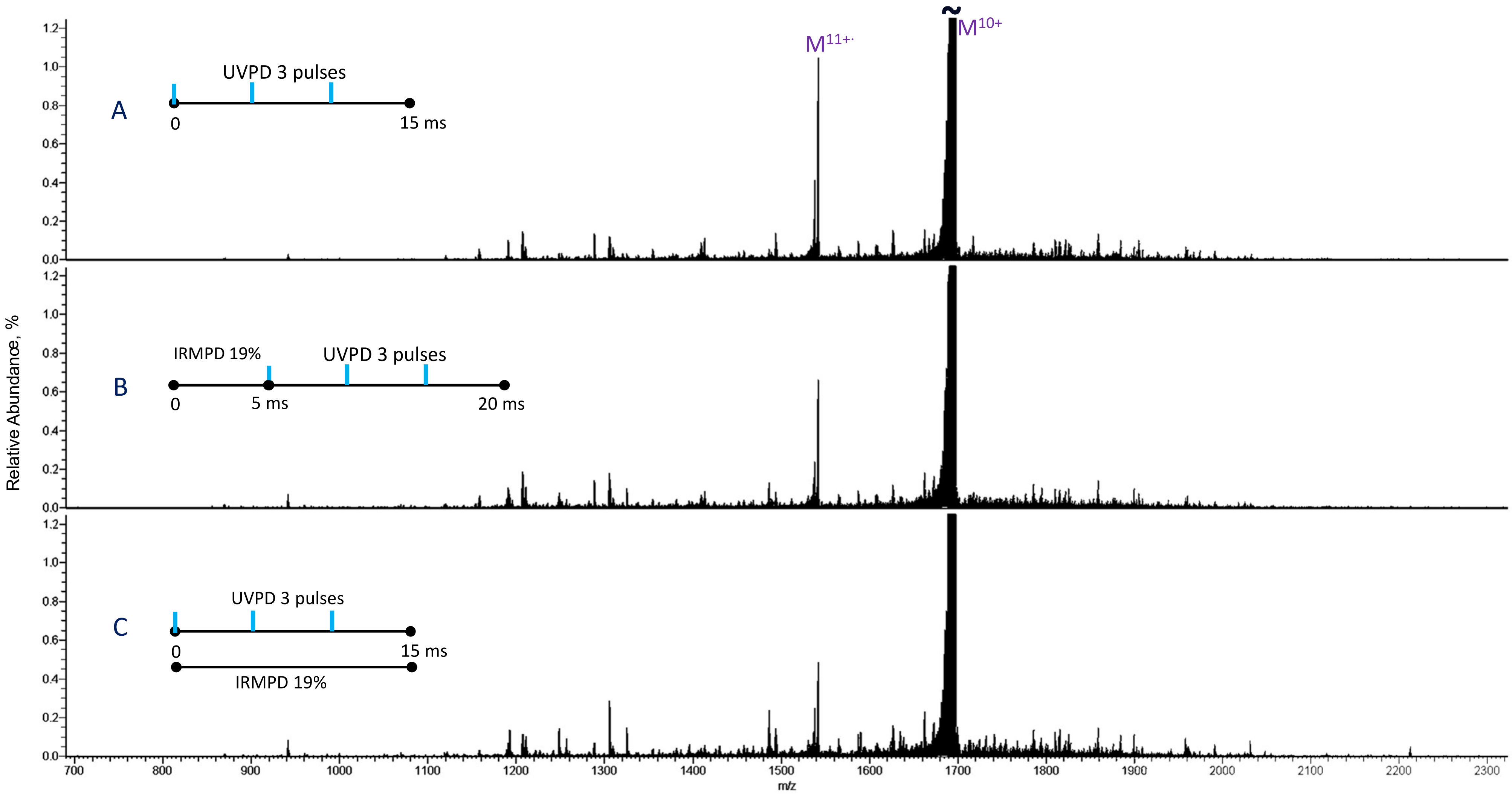


Figure S15. UVPD and IR-activated-UVPD spectra of myoglobin$^{10+}$. A) UVPD; B) IRMPD 5 ms (19 % laser duty cycle) followed by UVPD; C) continuous irradiation of precursor ions by IR (19 % laser duty cycle) for 15 ms during UVPD. In all experiments, three pulses of UV laser, 5 mJ/pulse were used for UVPD.

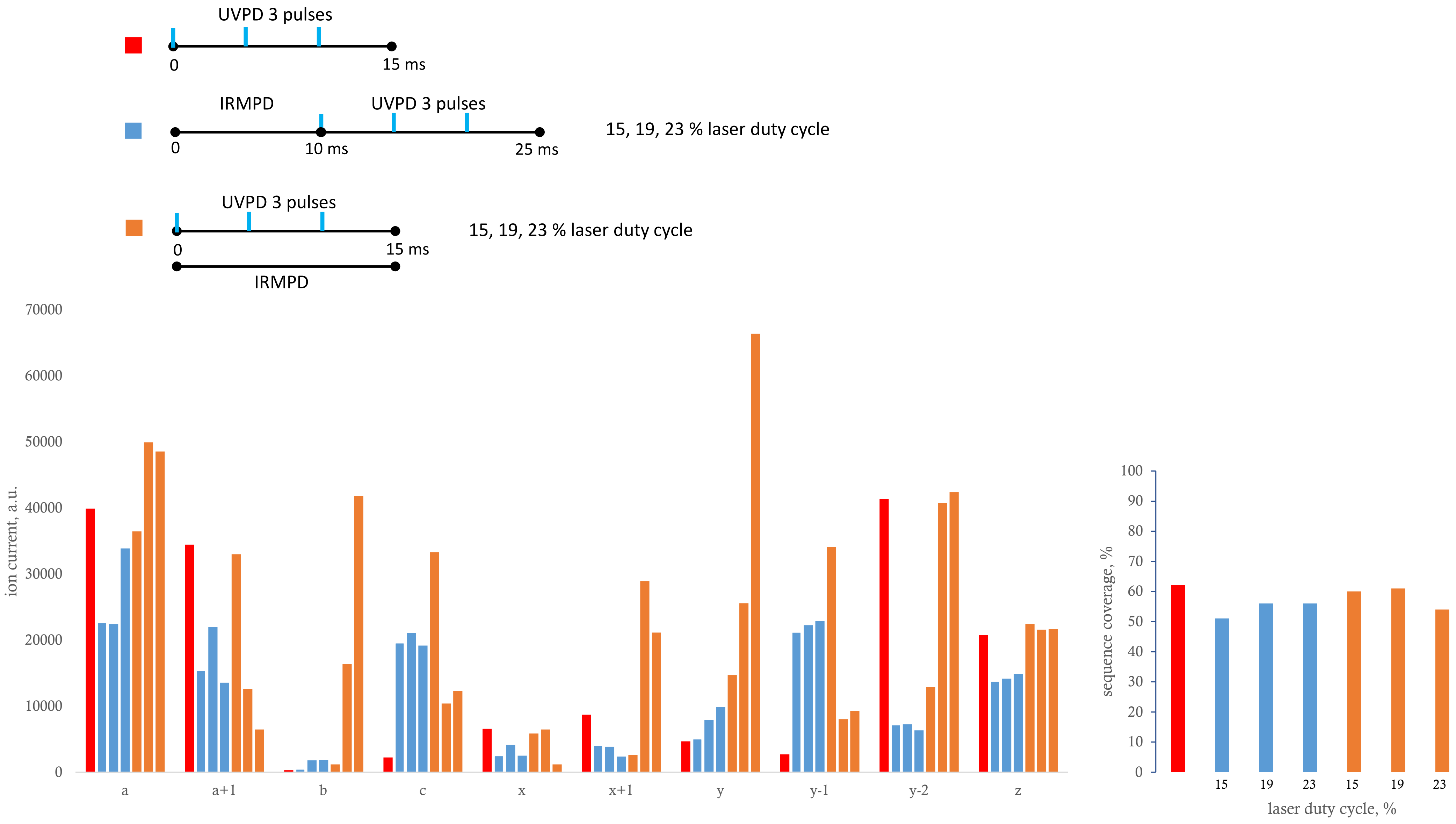


Fig S16. Ion currents of fragments of different types and sequence coverages identified in the IR-activated-UVPD experiments of myoglobin$^{10+}$.

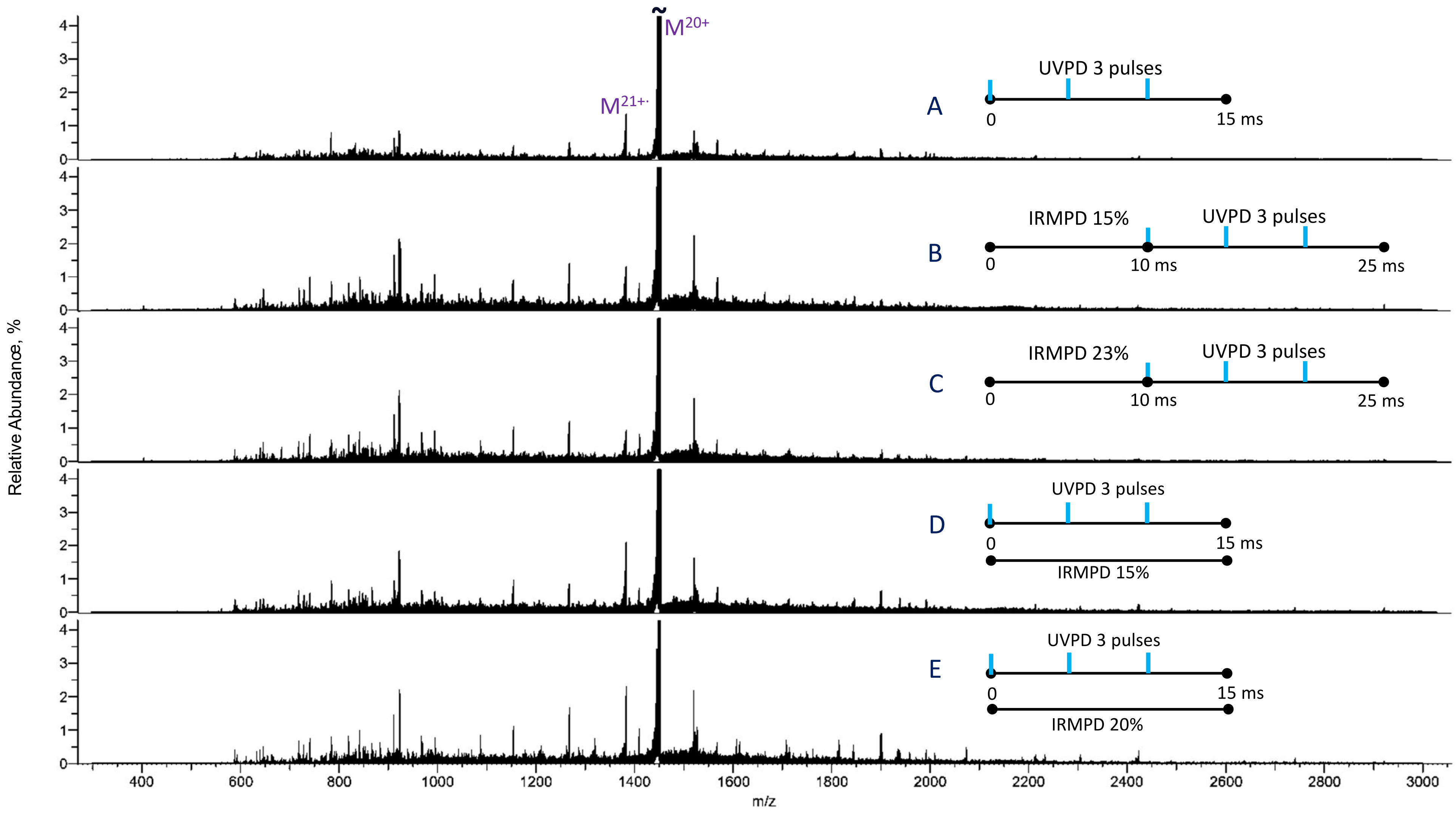


Figure S17. UVPD and IR-activated-UVPD spectra of [carbonic anhydrase]$^{20+}$. A) UVPD; B) IRMPD 10 ms (15 % laser duty cycle) followed by UVPD; C) IRMPD 10 ms (23 % laser duty cycle) followed by UVPD; D) continuous irradiation of precursor ions by IR (15 % laser duty cycle) for 15 ms during UVPD; E) continuous irradiation of precursor ions by IR (20 % laser duty cycle) for 15 ms during UVPD. In all experiments, three pulses of UV laser, 5 mJ/pulse were used for UVPD.

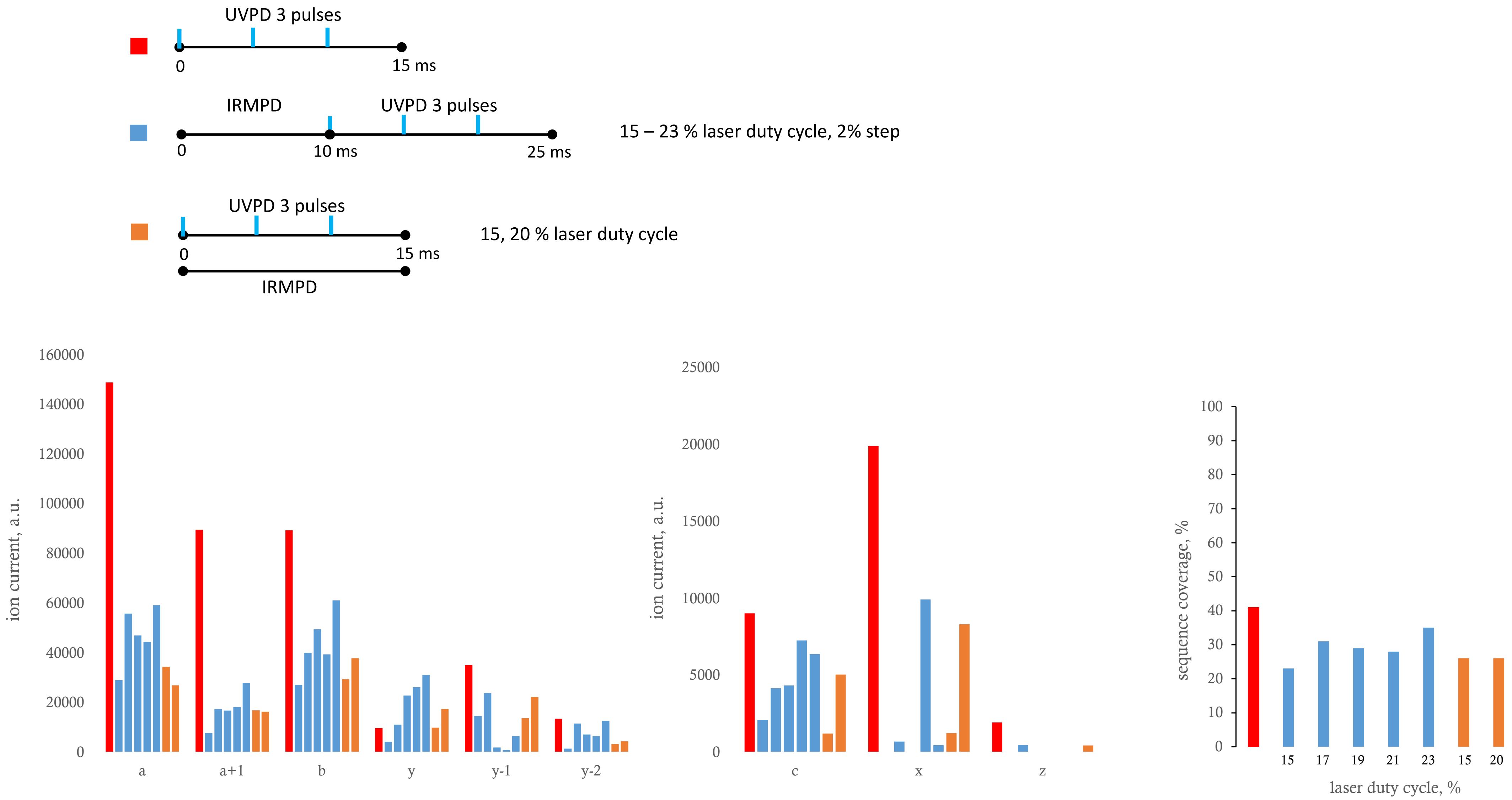


Fig S18. Ion currents of fragments of different types and sequence coverages identified in the IR-activated-UVPD experiments of [carbonic anhydrase]$^{20+}$.

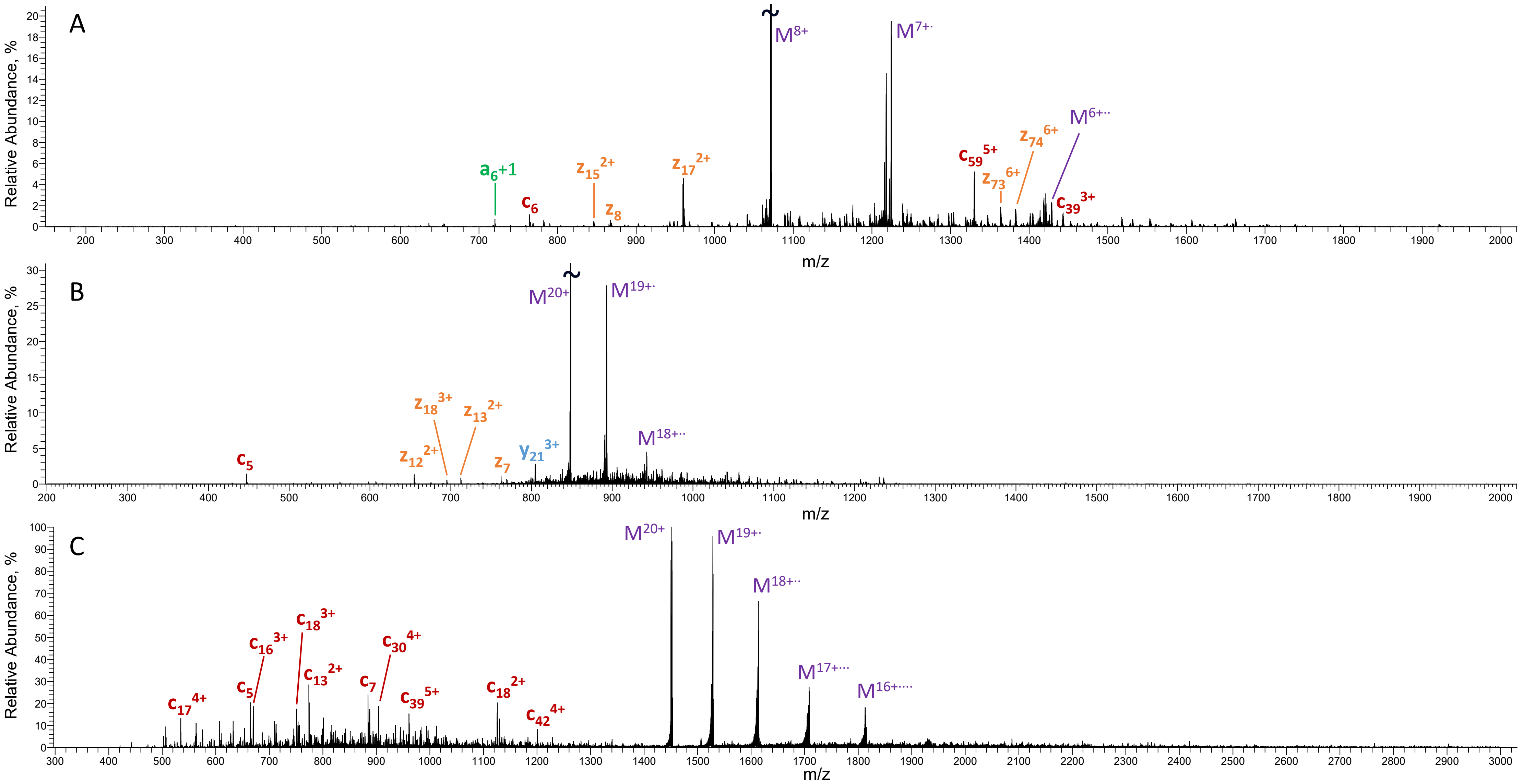


Fig S19. ECD spectra of ubiquitin$^{8+}$ (A), myoglobin$^{20+}$ (B), and [carbonic anhydrase]$^{20+}$ (C) acquired following 50 ms, 30 ms, and 20 ms of irradiation by 2 eV electrons, respectively. For clarity, only few selected fragments are annotated

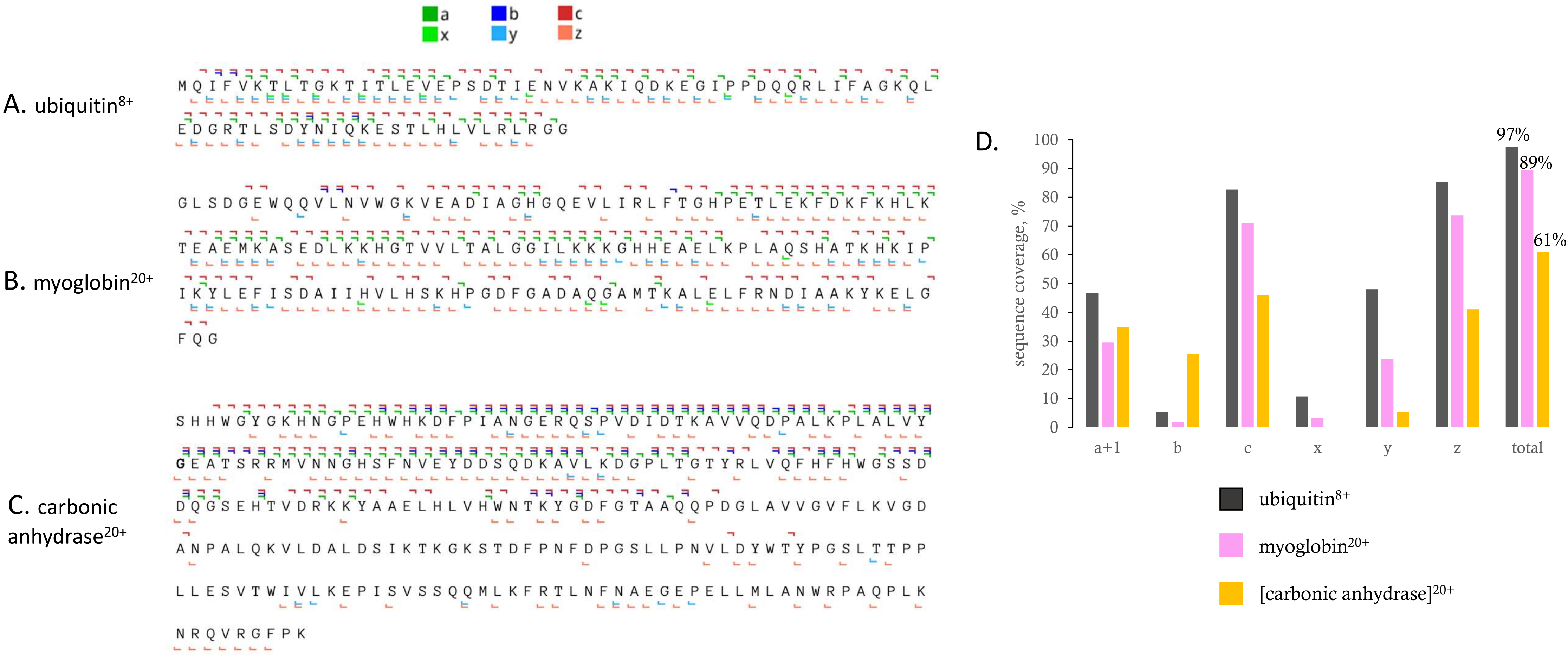


Figure S20. Fragment maps of ECD of A) ubiquitin$^{8+}$ (50 ms irradiation), B) myoglobin$^{20+}$ (30 ms irradiation), and C) [carbonic anhydrase]$^{20+}$ (20 ms irradiation). D) Sequence coverages by fragments of different types identified in the ECD experiments of ubiquitin$^{8+}$ (50 ms irradiation), myoglobin$^{20+}$ (30 ms irradiation), and [carbonic anhydrase]$^{20+}$ (20 ms irradiation).

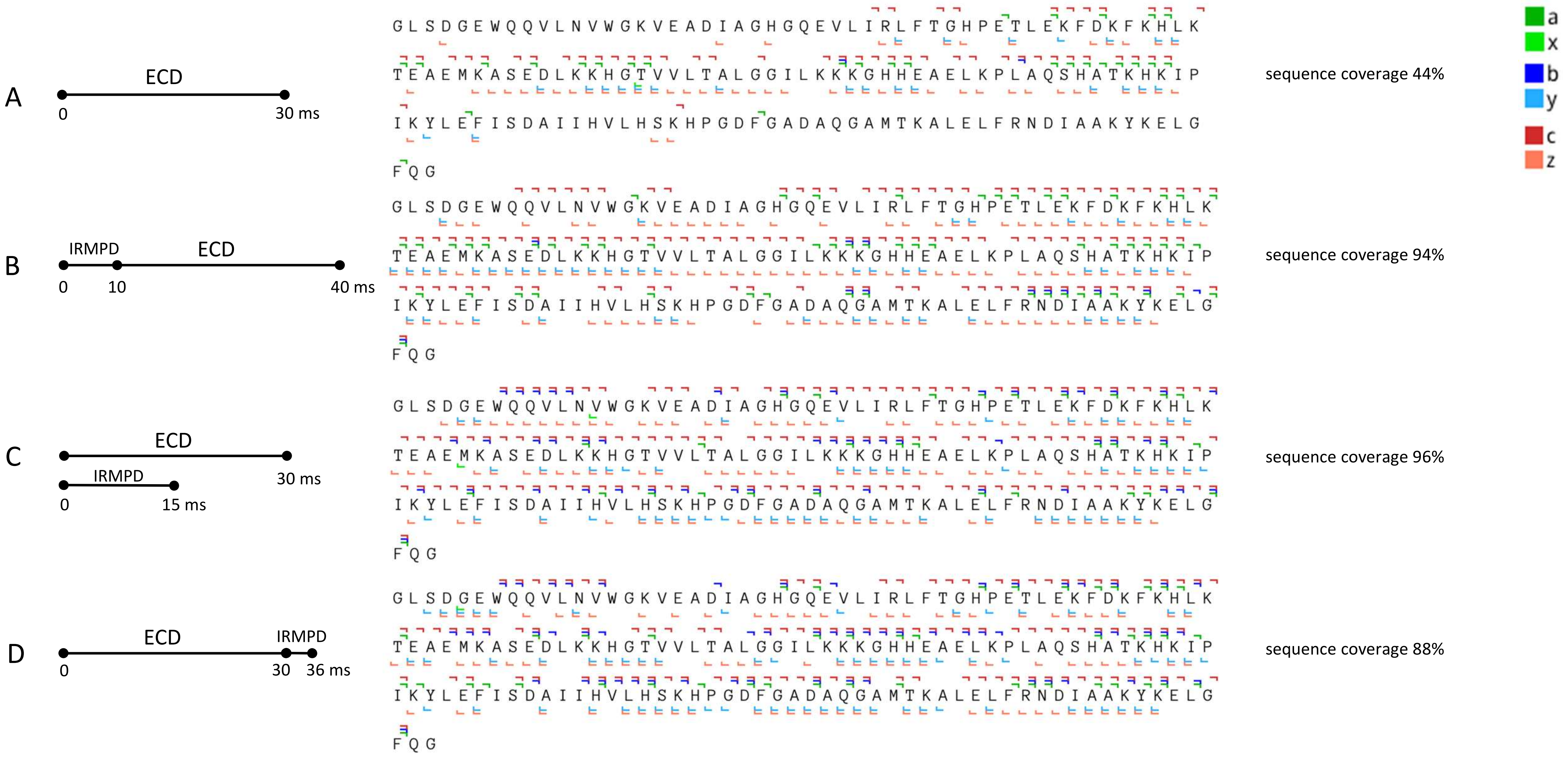


Figure S21. Fragment maps of ECD and IR-activated-ECD of myoglobin$^{10+}$ as given in Figure 4. A) ECD 30 ms; B) IRMPD 10 ms (15 % laser duty cycle) followed by 30 ms ECD; C) IRMPD (14 % laser duty cycle) concurrently with ECD for 15 ms followed by 15 ms of ECD; D) ECD 30 ms followed by 6 ms IRMPD (19 % laser duty cycle). The energy of electrons in all experiments was 2 eV.

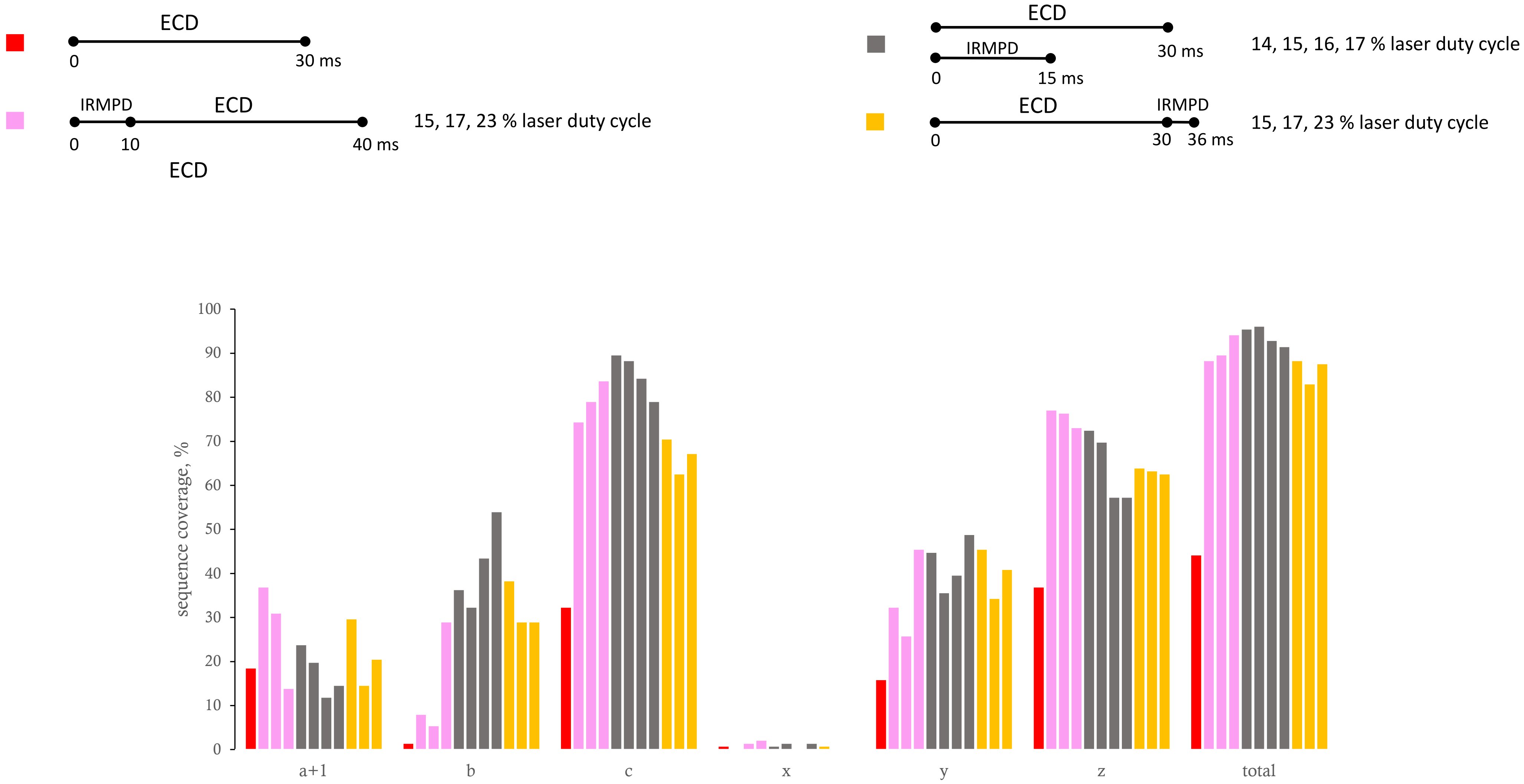


Fig S22. Sequence coverages by fragments of different types identified in the ECD and IR-activated-ECD experiments of myoglobin$^{10+}$ as given in Figure 4.

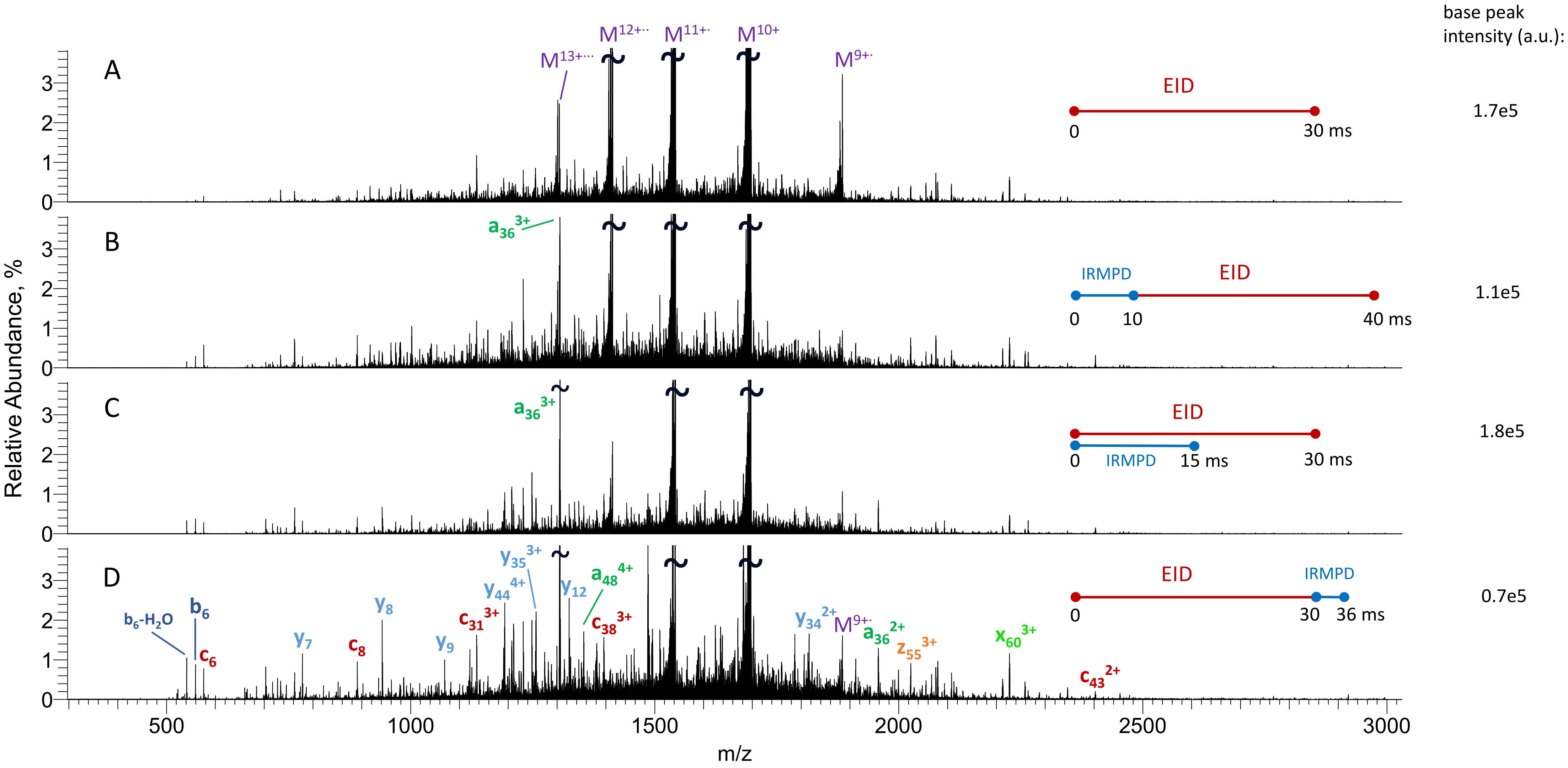


Figure S23. EID and IR-activated-EID spectra of myoglobin$^{10+}$. A) EID 30 ms; B) IRMPD 10 ms (15 % laser duty cycle) followed by 30 ms EID; C) simultaneous irradiation of precursor ions by IR light (14 % laser duty cycle) and electrons for 15 ms followed by 15 ms of EID; D) EID 30 ms followed by 6 ms IRMPD (19 % laser duty cycle). For clarity, only few selected fragments are annotated

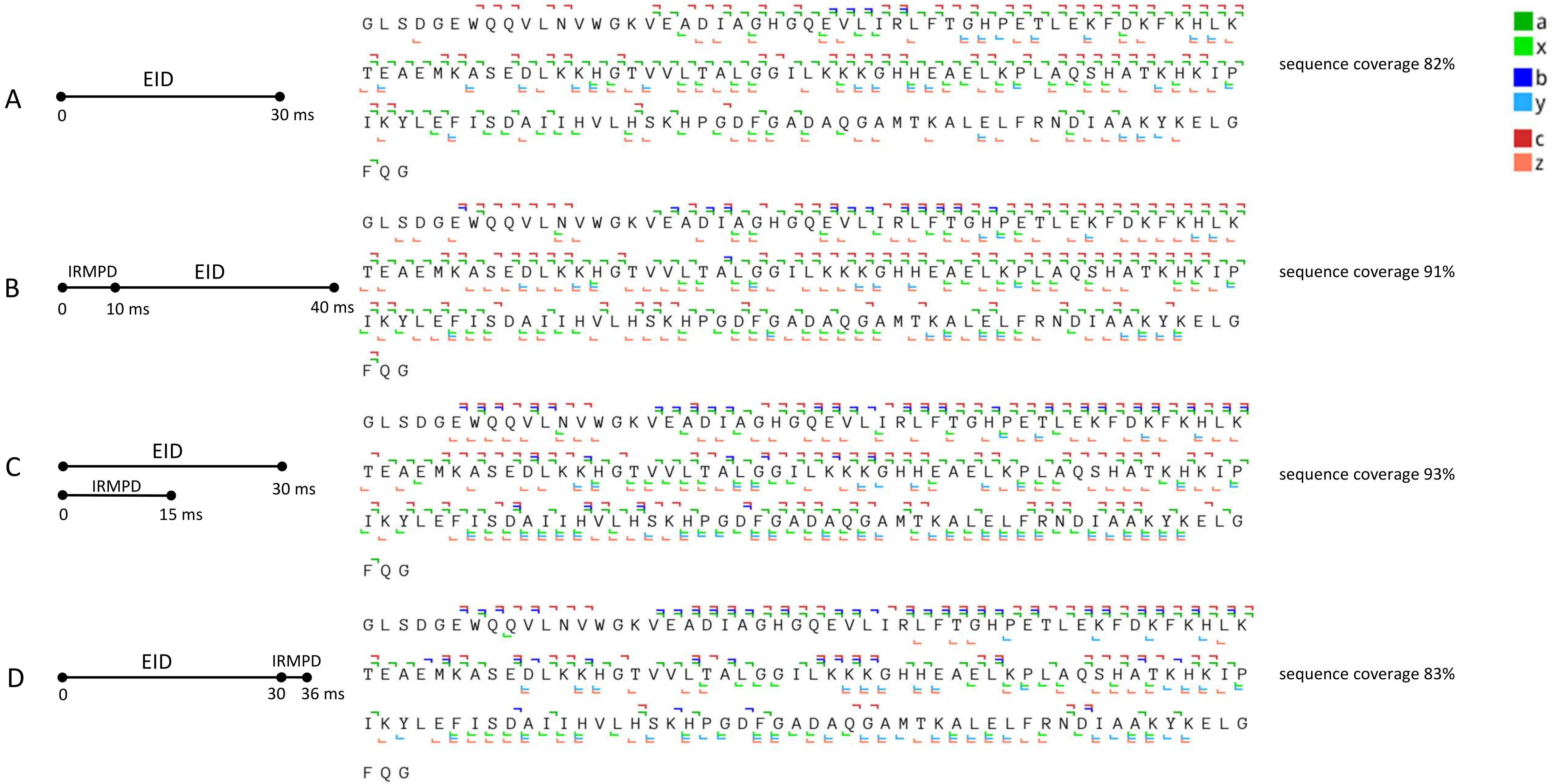


Figure S24. Fragment maps of EID and IR-activated-EID of myoglobin$^{10+}$. A) EID 30 ms; B) IRMPD 10 ms (15 % laser duty cycle) followed by 30 ms EID; C) simultaneous irradiation of precursor ions by IR light (14 % laser duty cycle) and electrons for 15 ms followed by 15 ms of EID; D) EID 30 ms followed by 6 ms IRMPD (19 % laser duty cycle). The energy of electrons in all experiments was 35 eV.

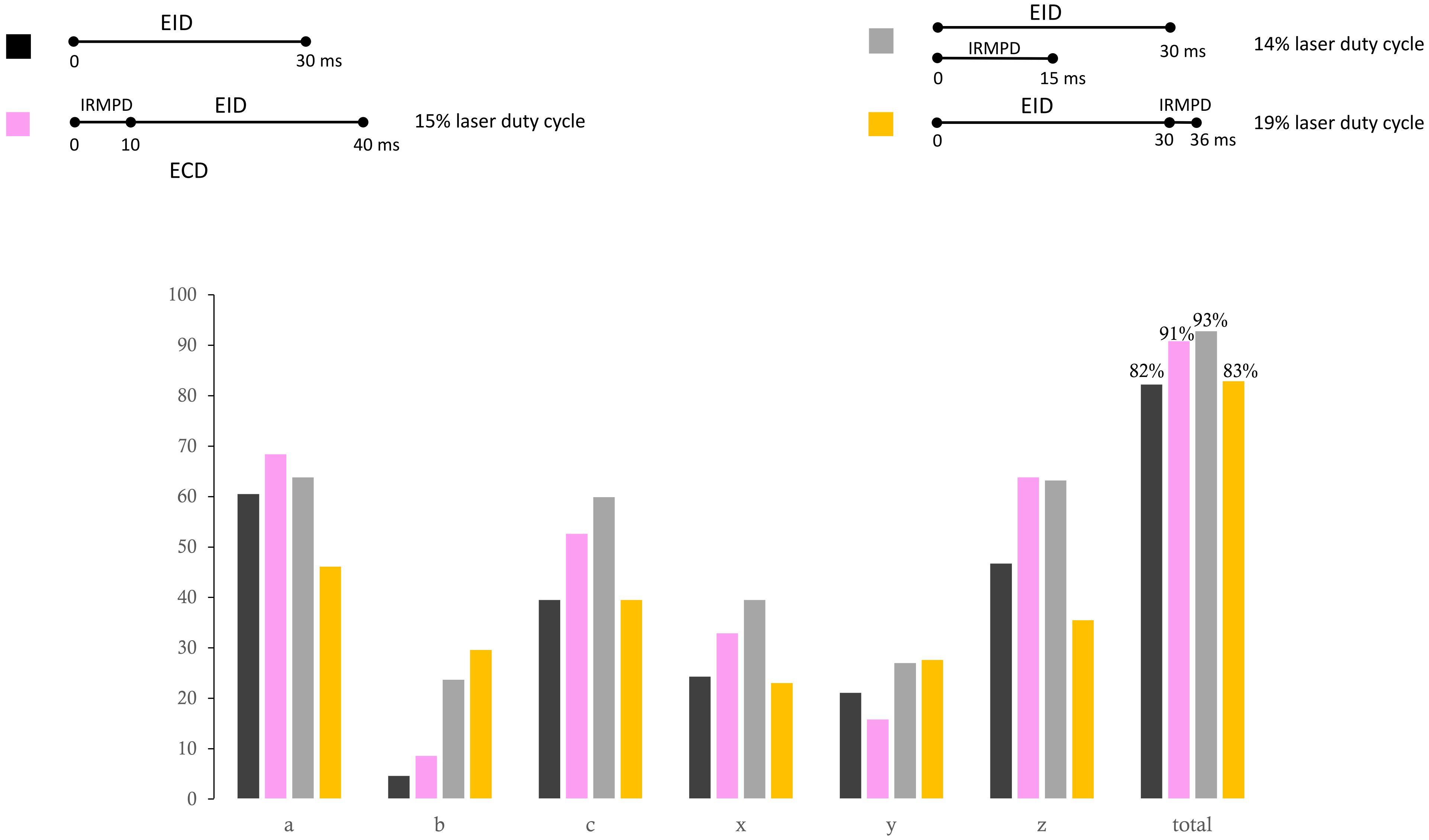


Fig S25. Sequence coverages by fragments of different types identified in the EID and IR-activated-EID experiments of myoglobin$^{10+}$ as given in Figure S23.

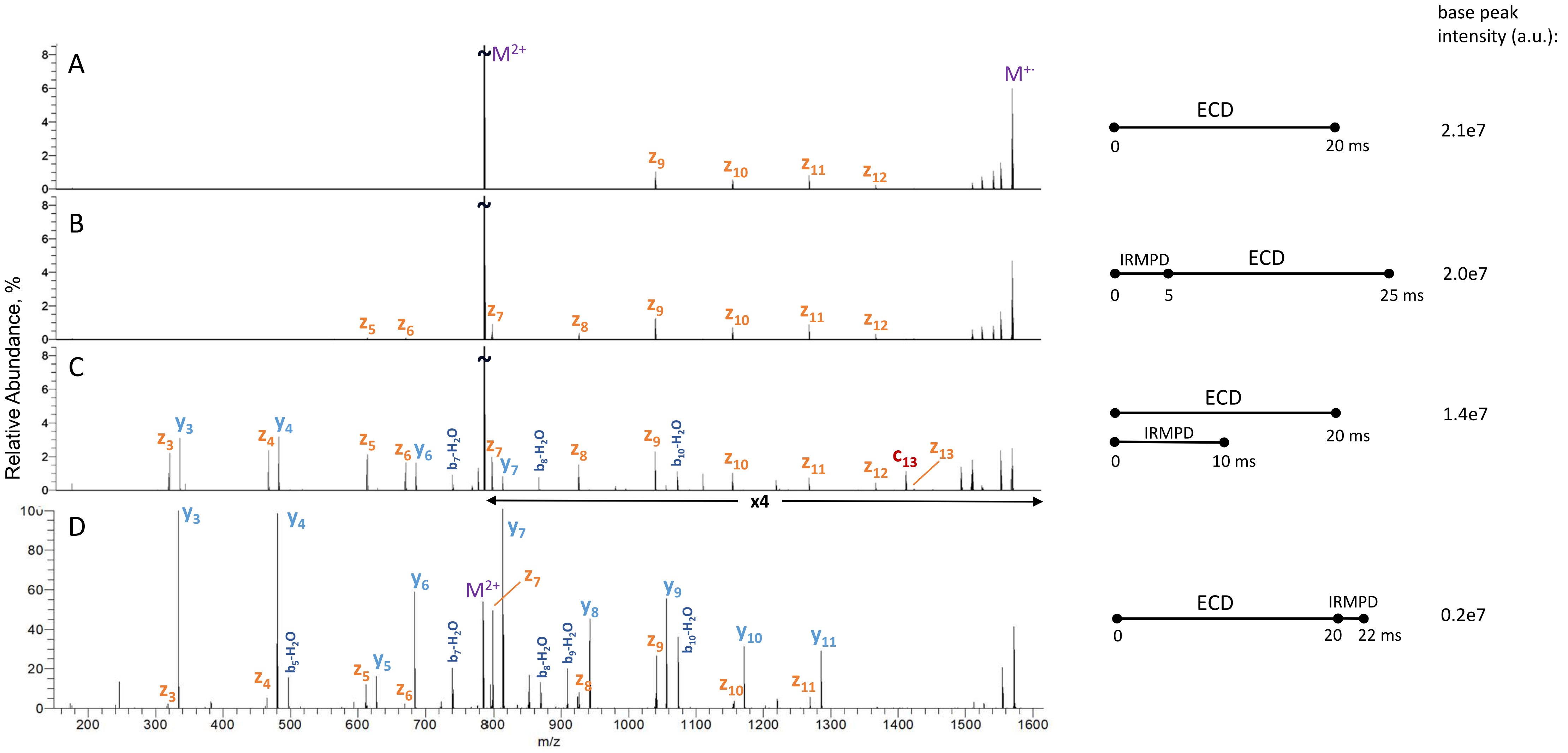


Figure S26. ECD and IR-activated-ECD spectra of [Glu-fibrinopeptide B]$^{2+}$. A) ECD 20 ms; B) IRMPD 5 ms (19 % laser duty cycle) followed by 20 ms ECD; C) simultaneous irradiation of precursor ions by IR (17 % laser duty cycle) and ECD for 10 ms followed by 10 ms of ECD; D) ECD 20 ms followed by 2 ms IRMPD (37 % laser duty cycle)

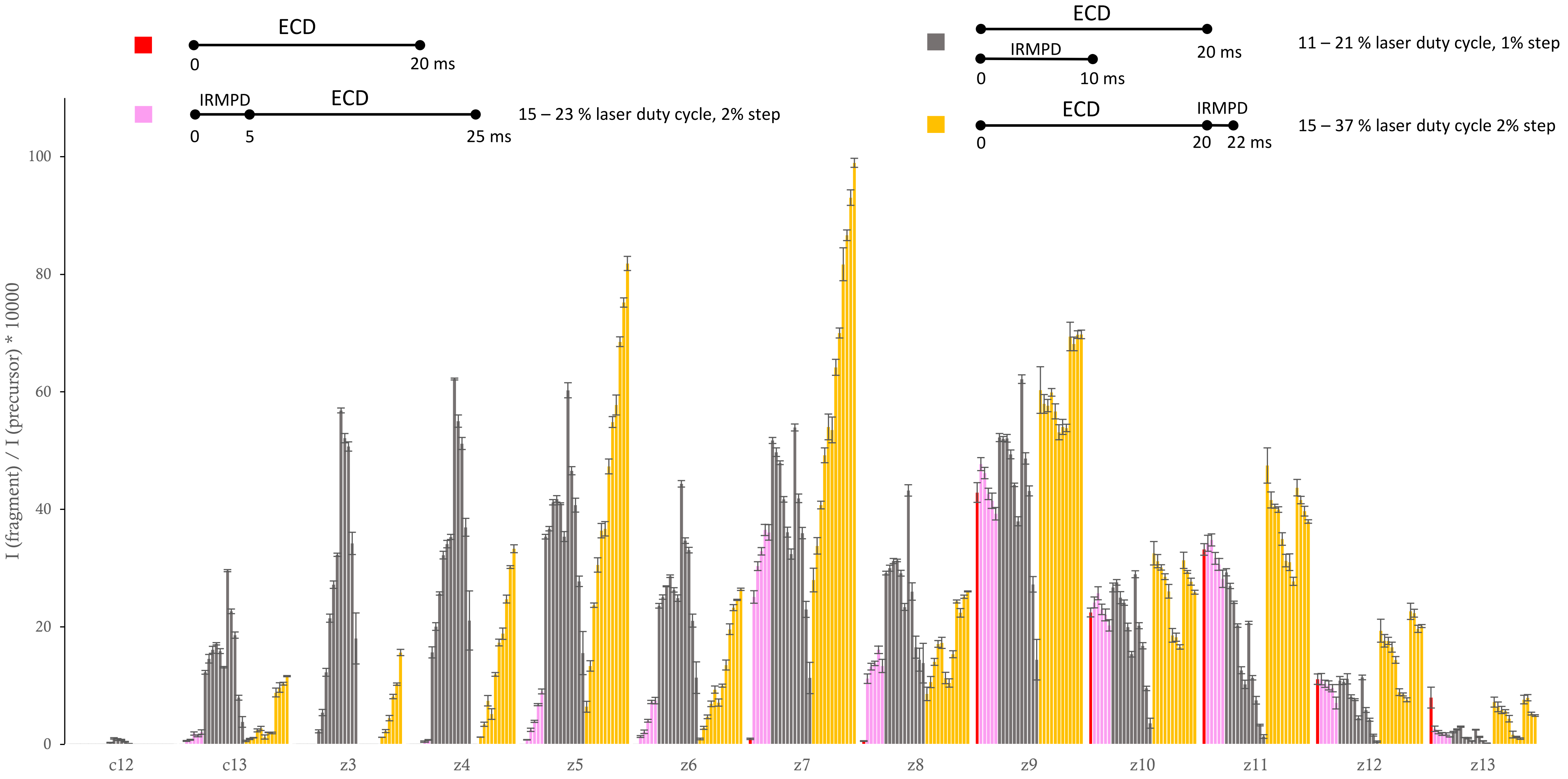


Figure S27. Distributions of normalised intensities of all *c* and *z* fragments identified in IR-activated-ECD of [Glu-fibrinopeptide B]$^{2+}$ as described in Figure S20 for different power outputs of the IR laser. The experiment was run in triplicate. Each data bar represents an average of a triplicate with standard deviations given as error bars. The intensities of fragments in each acquisition were normalised to the intensity of the isolated precursor in MS1.

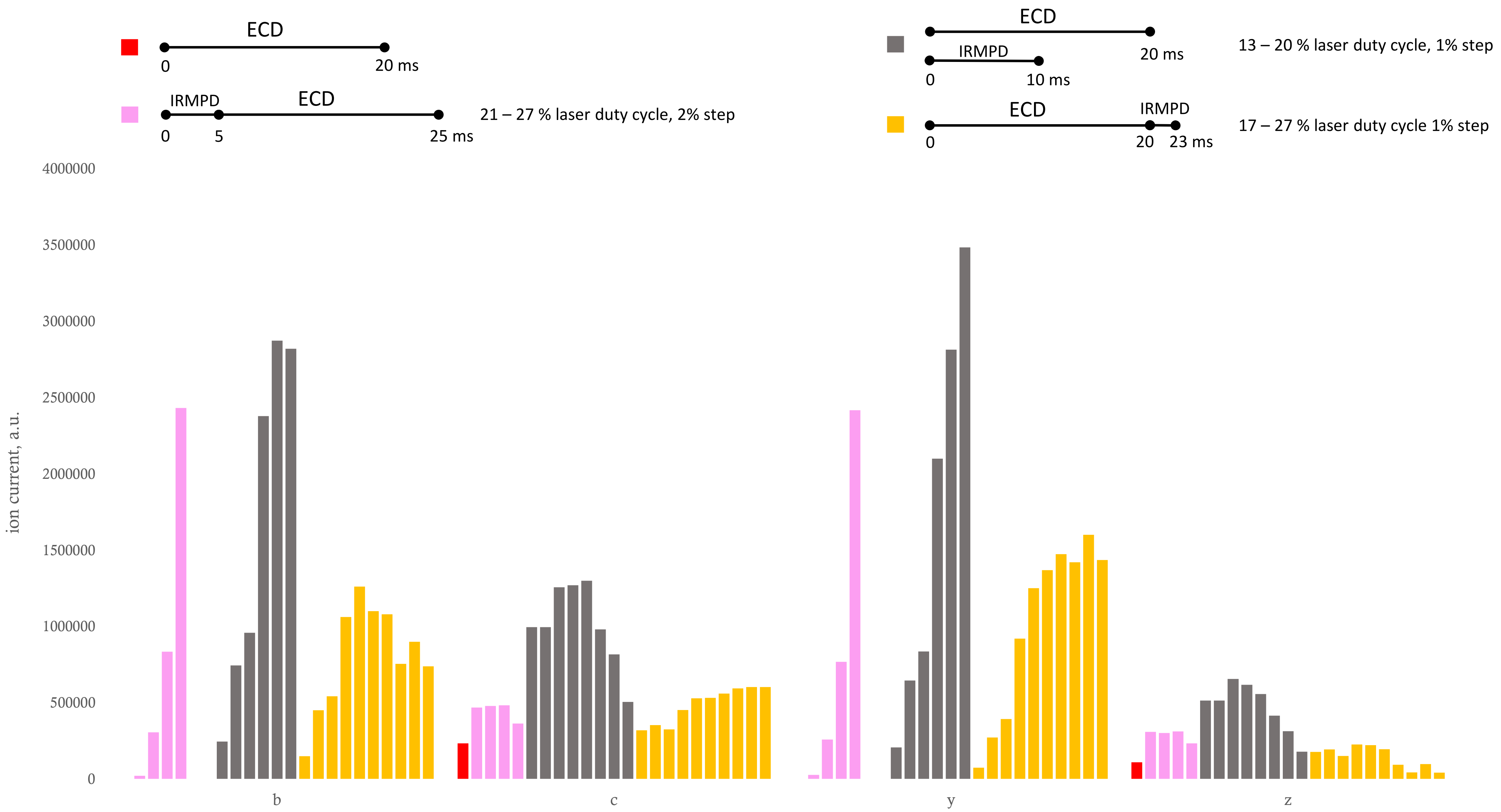


Figure S28. Summed ion currents of *a*,*b*,*c*,*y*,*z* fragments identified in IR-activated-ECD of triply charged chain B of insulin as described in Figure S22 for different power outputs of the IR laser.
